# Dysregulation of excitatory neural firing replicates physiological and functional changes in aging visual cortex

**DOI:** 10.1101/2020.01.06.896324

**Authors:** Seth Talyansky, Braden A. W. Brinkman

**Author notes:** Now at Stanford University (Seth Talyansky).

## Abstract

The mammalian visual system has been the focus of countless experimental and theoretical studies designed to elucidate principles of neural computation and sensory coding. Most theoretical work has focused on networks intended to reflect developing or mature neural circuitry, in both health and disease. Few computational studies have attempted to model changes that occur in neural circuitry as an organism ages non-pathologically. In this work we contribute to closing this gap, studying how physiological changes correlated with advanced age impact the computational performance of a spiking network model of primary visual cortex (V1). Our results demonstrate that deterioration of homeostatic regulation of excitatory firing, coupled with long-term synaptic plasticity, is a sufficient mechanism to reproduce features of observed physiological and functional changes in neural activity data, specifically declines in inhibition and in selectivity to oriented stimuli. This suggests a potential causality between dysregulation of neuron firing and age-induced changes in brain physiology and performance. While this does not rule out deeper underlying causes or other mechanisms that could give rise to these changes, our approach opens new avenues for exploring these underlying mechanisms in greater depth and making predictions for future experiments.

## Introduction

While healthy aging, and in particular its impact on neurological performance, has been the focus of numerous experimental studies [1–8], there are comparatively few theoretical and computational studies that focus on normal aging. Most computational studies to date model the effects of diseases that often manifest in advanced age in humans [9,10], such as Alzheimer’s [11–22] or Parkinson’s disease [23–30]. Only recently has theoretical and computational work on aging in non-pathological networks begun to emerge [31,32]. Because advanced age is one of the most important risk factors for developing such neurological disorders [9,10,33], to fully understand the progression of these diseases we ought to have a baseline understanding of how the brain’s circuitry changes during healthy aging, both in terms of physiological properties and functional performance. This would help dissociate disease-related changes from those caused during normal aging, and thereby allow researchers to focus their attention on treating potential causes of the disease progression not directly related to normal aging. On the other hand, understanding how the healthy brain ages may enable us to treat declines in performance caused solely by aging, in both healthy subjects and those with neurological disorders or diseases.

In this work we seek to advance our understanding of potential mechanisms and consequences of age-induced changes in physiology and performance in visual cortex. We do so by using a V1-like network model to qualitatively replicate experimentally-observed differences in the physiology and function of neurons in the visual cortex of young and old cats. In particular, we are motivated by experimental work that finds that senescent brain tissue in cat V1 shows increased spontaneous firing [34], decreased GABAergic inhibition [35,36], and decreased selectivity to the orientation of grating stimuli [34].

The logic of our approach is to mimic the experimental conditions by creating a V1-like spiking network model that we “age” in such a way that it develops the physiological changes observed in the experiments of [34–36]. Then, we can test the sensitivity of neurons to oriented grating stimuli at each age of our network to observe how neural selectivity changes as the networks become more and more senescent. Importantly, because we model the aging process, rather than constructing two discrete “young” and “old” networks, we will not only qualitatively replicate the experimental findings in young and elderly cats, but will also probe the model at intermediate ages to understand changes in physiology and function throughout aging.

In order to model the experimental results we use E-I Net, a previously-developed spiking network model of mammalian V1 [37], as the basis for our study. The network structure of E-I Net is learned by training the model on pixel images of natural scenes, mimicking the early developmental processes [38] in which neurons develop Gabor-like receptive fields measured experimentally in the visual cortex of mammals [39]. Similarly, the recurrent connections develop such that neurons with similar receptive fields effectively inhibit one another through intermediary inhibitory neurons. The standard E-I Net model thus contains the important V1-like features we seek in a network model, and we use it to represent the young, mature network. In order to “age” the network we modify and extend the training procedure to create the aging-like process that qualitatively reproduces the experimental results of [34,35], in addition to making predictions for other features of the aging process. Importantly, we show that the only modification we need to make to obtain a process that results in aged network phenotypes is a change to the training procedure that will promote increased firing activity within the network with age—the other physiological and performance changes emerge from this single change. We argue that this change can be interpreted as a breakdown in homeostatic-like mechanisms controlling the network’s target firing rate, leading to the increases in excitation, decreases in synaptic strength, and declines in orientation selectivity that mimic the experimentally observed changes. Although there may be other mechanisms that can replicate these experimental findings, our model makes several predictions for other yet-to-be-observed changes in physiology and functional performance that can potentially be used to further test our model and the interpretation of our results.

This paper proceeds as follows: in Results we first give a brief overview of E-I Net and the modifications we made to the model to implement an aging-like phase of the training dynamics. We then present our findings on the physiological changes that occurred in our network during this aging phase—i.e., the changes to receptive fields, lateral connections, and firing thresholds—followed by analyses of the changes in functional performance of the network, which we quantify by the changes in the selectivity of the neurons to oriented gratings. To gain a deeper understanding of how each set of our network parameters contributes to performance changes during the aging process, we perform a number of numerical experiments. In particular, i) we study how orientation selectivity evolves when one or more parameters sets are frozen to their young values and ii) we disambiguate whether receptive field structure or receptive field magnitude contributes more to the declines in orientation selectivity. Finally, in the Discussion we interpret our results and their relationship to recent work, the limitations of our approach, how our model’s predictions might be tested experimentally, and possible implications for understanding normal and pathological declines in neural performance with age.

## Results

### Modified E-I Net as a model of aging in V1-like networks

We begin by briefly reviewing E-I Net [37], the spiking network model of visual cortex we use in this work, and outlining how we modify this model to implement the aging-like process we use to replicate the experimental findings of [34,35]. E-I Net comprises a population of recurrently-connected excitatory and inhibitory spiking neurons that receive external inputs driven by visual stimuli. The subthreshold dynamics of the neurons’ membrane potentials obey

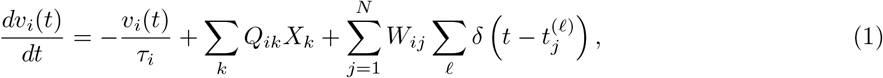

where *v_i_*(*t*) is the membrane potential of neuron *i* at time *t* and the membrane time constant *τ_i_* is *τ_E_* (*τ_I_*) if neuron i is excitatory (inhibitory). Neuron *i* fires a spike when *v_i_*(*t*) = *θ_i_* and then is reset to *v_i_*(*t*) = 0. As shown schematically in Fig. 1, each of the *N* neurons receives visual input in the form of pixel images *X* from natural scenes. The input currents generated by these visual scenes appearing in Eq. (1) is ∑_*k*_ *Q_ik_X_k_*, where *X_k_* is the intensity of the *k*^th^ pixel of the input patch *X* and the *Q_ik_* are the strengths of the input connections from the *k*^th^ pixel to neuron *i*. Neurons are connected by recurrent synaptic connections *W_ij_*, representing the amount of charge transferred when pre-synaptic neuron *j* transmits a spike to post-synaptic neuron *i*. While the individual *W_ij_* are distinct for all pairs *ij,* there are three classes of recurrent weights: inhibitory-to-excitatory weights *W_EI_*, excitatory-to-inhibitory weights *W_IE_*, and inhibitory-to-inhibitory weights *W_II_*; as in the original E-I Net model we assume excitatory-to-excitatory weights *W_EE_* are rare and may be neglected. The time at which the ℓ^th^ spike is emitted by neuron *j* is represented by 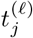 in Eq. (1). In this paper we take *W_ij_* to be positive (negative) if the post-synaptic neuron j is excitatory (inhibitory), in contrast to the original E-I Net paper which makes the sign explicit, such that *W_ij_* is non-negative for any cell type.

**Figure 1:**
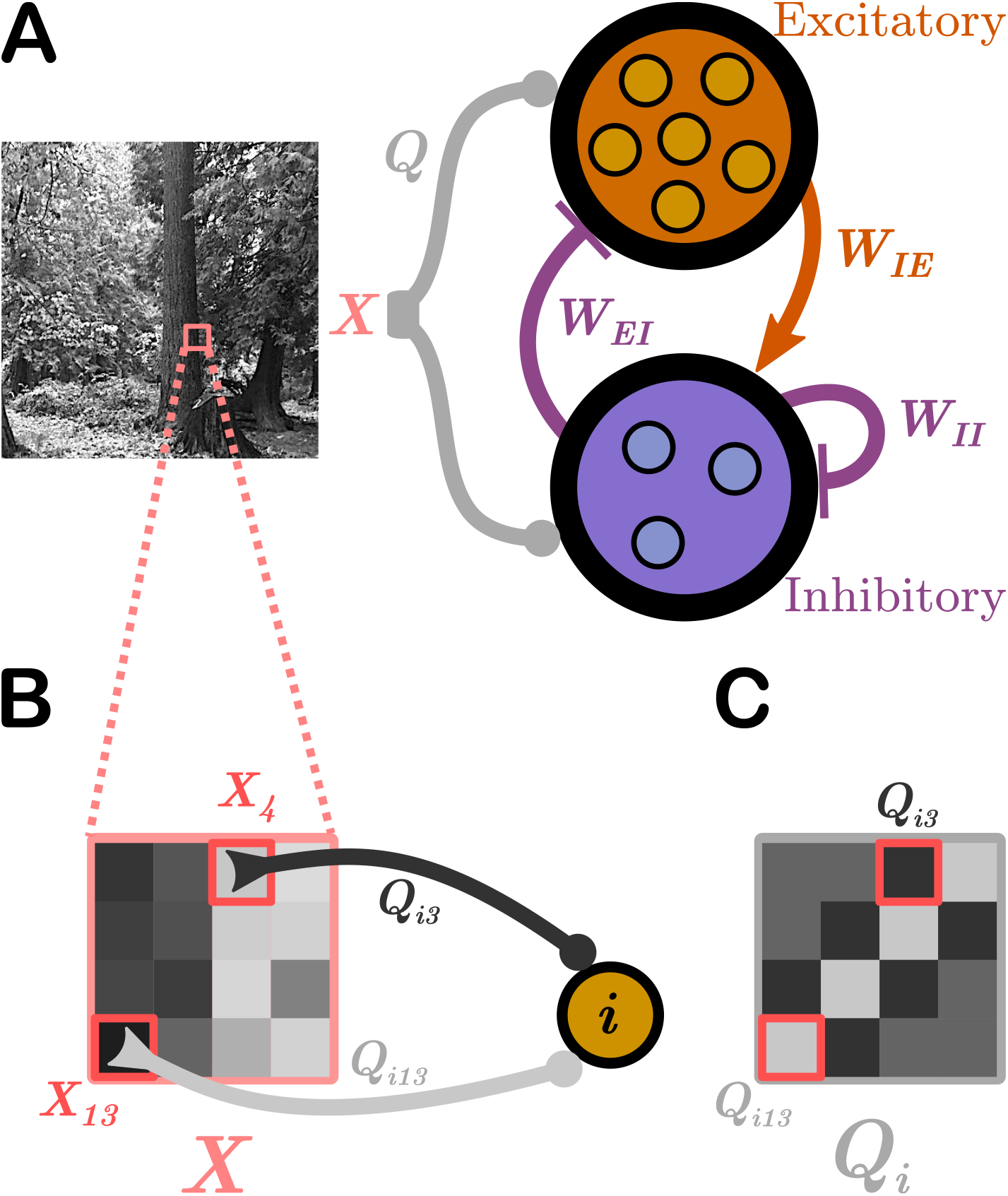
Schematic of the network model: We use E-I Net [37] as our model of visual cortex. **A**. E-I Net comprises a network of leaky integrate-and-fire neurons that receive visual input in the form of randomly-drawn patches from pixel images (*X*, highlighted region of natural scene), communicated to the neurons through input weights *Q.* Each neuron belongs to either the excitatory or inhibitory population. Neurons are synaptically connected by lateral weights *W* that obey Dale’s law, such that excitatory-to-inhibitory weights *W_IE_* are positive, while inhibitory-to-excitatory (*W_EI_*) and inhibitory-to-inhibitory (*W_II_*) weights are negative. Excitatory-to-excitatory connections are assumed to be negligible and are not included in this model. The input weights *Q*, lateral weights *W*, and firing threshold of each neuron *θ* (not depicted) are learned by training the network on natural pixel images. **B**. The input stimulus to the network, *X*, comprises a grid of pixels (4 × 4 shown here; 8 × 8 used in practice). The *k*^th^ pixel generates a current in the *i^th^* neuron equal to the intensity of the pixel multiplied by a weight *Q_ik_*, for a total current into neuron *i* of ∑_k_ *Q_ik_X_k_*. **C**. The input weights *Q* can be arranged spatially by ordering them the same way as the pixel-input patches. This representation can be interpreted as neuron *i*’s receptive field.

These parameters develop according to synaptic and homeostatic learning rules inspired by sparse coding theory and previously established rules such as Oja’s Hebbian plasticity rule [40] and Foldiak’s rule [41]. These learning rules are not specific to any particular species, and therefore the network is taken to be a general model of mammalian visual cortex. The learning rules are designed to enforce linear decoding of the image patches from the spike counts (Eq. (2)), minimal correlation in spike counts (Eq. (3)), and sparseness in firing (Eq. (4)):

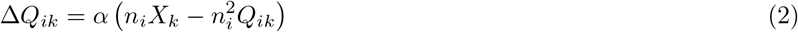

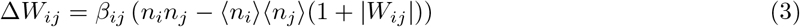

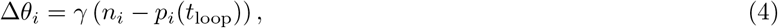

where *n_i_* is the spike count of neuron i in response to the presented image patch over a duration *T*, 〈*n_i_*〉 is a moving average over time, weighted by an exponential decay factor, and the parameters *α*, *β_ij_*, and *γ* are the learning rates, which set how quickly the network parameter updates converge to a steady state. The synaptic learning rates *β_ij_* ∈ {*β_EI_*, *β_IE_*, *β_II_*} depend on whether the types of neurons connected by the synapse (excitatory or inhibitory). The quantity *p_i_*(*t*_loop_) is the desired mean number of spikes emitted per time step, evaluated at the “age” of the network *t*_loop_, quantified by the number of training loops that have taken place (i.e., the number of updates to the network parameters). Sparsity in firing is enforced by choosing *p_i_*(*t*_loop_) ≪ 1. In the original E-I Net model *p_i_*(*t*_loop_) is a constant value *p_E_*(*p_I_*) if neuron *i* is excitatory (inhibitory), independent of the network age.

In the original E-I Net model the learned network parameters *Q_ik_,W_ij_*, and *θ_i_* achieve a steady-state network structure in which the pixel-space arrangements of input weights *Q* resemble Gabor-like receptive fields typical of networks trained on natural scenes [39], as shown schematically in Fig. 1C. The learned lateral weights are organized such that excitatory neurons with similar receptive fields effectively inhibit one another [39,42]. To study the effect of aging, and in particular how the network structure changes with age, we modify the target firing rate *p_E_*(*t*_loop_) in Eq. (4), choosing it to be

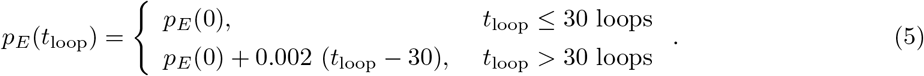

That is, after the network has achieved its steady state (which occurs by 30 training loops) the target spike count begins to increase with each loop. This change is designed to increase the spiking activity of the excitatory neurons in order to mimic the increased activity of neurons in aged cat brain tissue observed in [34]. Physiologically, this change may be interpreted as arising from a breakdown in homeostatic mechanisms that regulate the target spike rate, which we do not model explicitly (see Discussion).

As *p_E_*(*t*_loop_) increases, the network parameters will begin to depart from the previously stabilized values, and we can track their evolution with training time, or “age.” Moreover, at each of these ages we can assess both the physiological changes in *Q_ik_*, *W_ij_*, and *θ_i_*, as well as the functional performance of the network in response to different tasks. As outlined in the introduction, we seek to mimic the experimental conditions of [34], which measured the selectivity of young and elderly cat neurons to oriented grating stimuli and observed decreased selectivity in elderly cats. We therefore present patches of visual input from oriented grating images to our “aged” network models and quantify the selectivity of neurons to these edges.

We train our networks for an additional 50 loops, giving a total network age of 80 loops. The target spike rate *p_E_*(*t*_loop_) increases on every loop, and we record the aged set of parameters {*Q,W,θ*} and test the sensitivity to gratings every 5 loops. We report on the physiological changes during the aging process next (Physiological parameter changes), followed by the performance changes during the oriented grating tasks (Performance changes), and finally an additional set of numerical experiments that we ran to further probe how the different network parameters contributed to the observed changes in physiology and performance (Numerical experiments to test contribution of different physiological parameters to declines in performance).

### Physiological parameter changes

Fig. 2 displays the distributions of the learned physiological parameters in “(mature) youth” (30 training loops) compared to “old age” (80 training loops). We find the range of magnitudes of the synaptic and input weights decreases with age, while the firing thresholds expand by orders of magnitude. Note that the network model is formulated in dimensionless units, so the parameter ranges do not correspond directly to dimensionful quantities measured in experiments.

**Figure 2:**
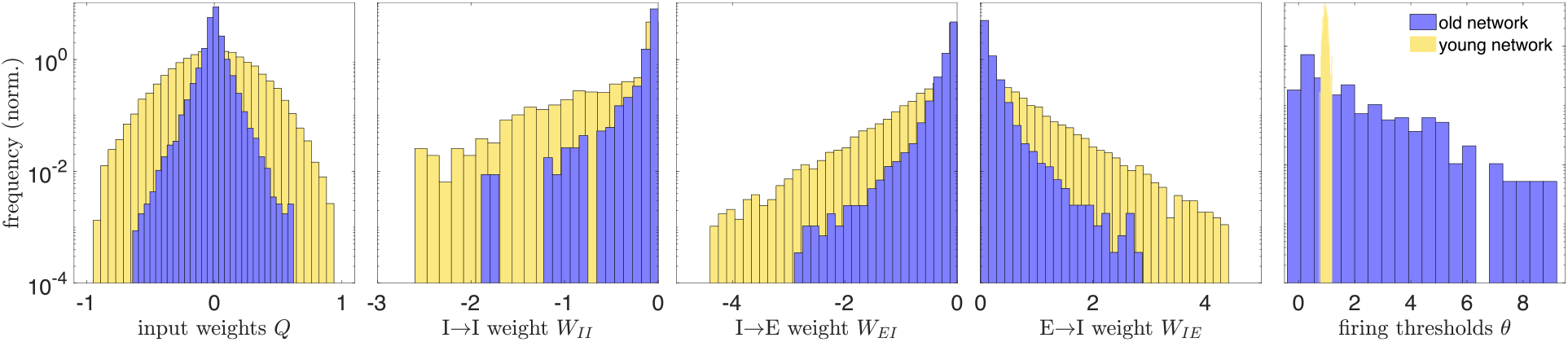
Histograms of physiological network parameters in youth vs. old age: Histograms of the magnitudes of network parameters learned during training on natural images: the input weights *Q*, lateral weights *W*, and firing thresholds *θ*. Yellow histograms correspond to the distributions of these parameters in the mature young network, the steady state of network development (reached by 30 training loops). After this age, the excitatory target firing rate begins to increase, causing a rearrangement of the network that we interpret as an aging-like process. The blue histograms are the distributions of network parameters after 80 training loops. Aging tends to decrease the magnitude of input and lateral weights, while it widens the distribution of firing thresholds (in young age, all the thresholds belong to a narrow range, relative to the old-age thresholds, and the bins are accordingly very narrow). A decrease in inhibitory weights is consistent with experimental work of [35, 36] and decreases in excitatory weights are consistent with experimental studies across several brain areas and species [5,6,8]. All quantities are expressed in dimensionless units.

In addition to changes in magnitude, the input weights also undergo a change in their spatial organization in pixel space—i.e., the receptive field of the neuron is not preserved with aging. Fig. 3A shows the receptive field of a particular neuron with a Gabor-like receptive field (RF) at age 30 (top) and a random-looking RF at age 80 (bottom). If we “vectorize” the receptive fields by rearranging the 8 × 8 pixel grids into 1 × 64 vectors, then we can quantify the “distance” between young and old receptive fields in pixel-space by computing the angular separation *φ_i_* between the vectorized young and older receptive fields *Q*^young^ and *Q*^old^ for each neuron *i*:

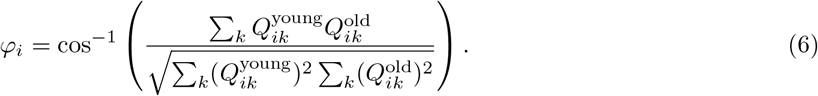

**Figure 3:**
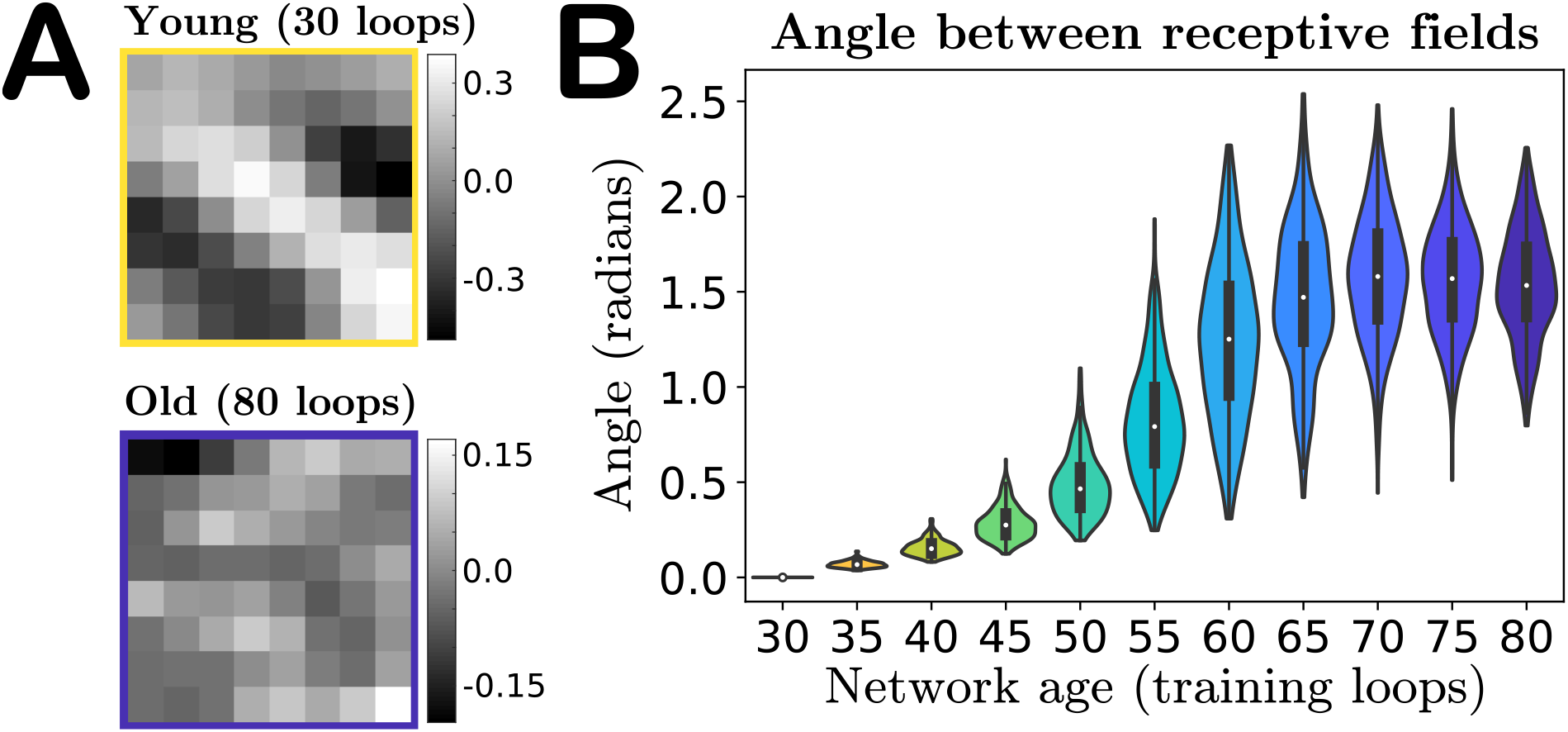
Angle between receptive fields in the input weight parameter space: **A**. A particular neuron’s receptive field (RF) in youth versus old age, demonstrating that features of a neuron’s RF may not be preserved in our model’s aging process. **B**. Violin plot of the distribution of angles between each individual excitatory neuron’s young and older vectorized receptive fields (RFs) (Eq. 6). As our network ages a neuron’s RF becomes increasingly orthogonal to its mature young RF, on average, indicated by the median of the RF distributions tending to approximately *π*/2. The colors of the violins represent the age indicated on the horizontal axis.

This angle is 0 if the old RF is identical to the young and *π*/2 if the young and old RFs are orthogonal. We find that the mean angle between vectorized RFs steadily increases with age, to the point that in the oldest age of our network the mean angle between a neuron’s young and old receptive fields tends to π/2, i.e., on average the old-age RFs of a population of neurons become orthogonal to the RFs in youth. Orthogonality in high-dimensions can be difficult to interpret, as random vectors in high-dimensional spaces are nearly orthogonal with high probability [43]. To gain some intuition into this result, we also compute the angles between the young (age 30) vectorized RFs and a shuffling of this same set of RFs, and compute the angles of separation between random pairs of these RFs. We obtain a distribution of angles (not shown) that is statistically indistinguishable from the distribution of angles between the young receptive fields and their age 80 counterparts (final violin plot in Fig. 3), using a two-sample Kolmogorov–Smirnov test (*p*-value of 0.36, failure to reject the null hypothesis that the empirical cumulative distribution functions come from the same distribution).

This result suggests that the aging process could simply be shuffling the RFs over time: no individual neuron’s receptive field is preserved, but across the network the distribution does not change significantly. To show that this is not the case, we classify the receptive fields of excitatory neurons as either “Gabor-like,” showing characteristics of Gabor edge detectors, or “not-Gabor,” and show that the numbers of neurons in these categories change with age. The results are given in Table 1. As some RFs categorized as not-Gabor in youth become Gabor-like with age, we also record transitions between types (bottom subtable in Table 1). We assess Gabor-ness by fitting a spatial Gabor profile to each neuron’s receptive field and classify it as Gabor-like if the goodness-of-fit relative to the RF magnitude is sufficiently small; see Methods and [39]. We use a conservative goodness-of-fit criterion; nonetheless, under this criterion close to a third of the excitatory RFs are classified as Gabor-like in youth, while this fraction drops to nearly 1-in-20 in our oldest network, demonstrating that the aged networks undergo a net loss of neurons with Gabor-like RFs. Given the edge-detecting properties of Gabor-like receptive fields, we thus anticipate this loss plays a prominent role in the degradation of the sensitivity of the network to oriented grating stimuli, discussed in the next section.

**Table 1:** Quantification of Gabor-like receptive fields: We quantify whether each excitatory neuron in our network model has a Gabor-like receptive field or not, in both our mature young network (30 loops) and the oldest network (80 loops). We assess Gabor-ness by fitting a spatial Gabor profile to each neuron’s receptive field and classify it as Gabor-like if the goodness-of-fit relative to the RF magnitude is sufficiently small; see Methods and [39]. In the bottom subtable, the numerator in each fraction corresponds to classification in old age and the denominator corresponds to classification in youth. For example, of the 123 excitatory neurons with Gabor-like RFs in youth, only 6 retain Gabor-like RFs in old age, while of the 400 — 123 = 277 neurons that are not Gabor-like (“not-Gabor”) in youth we find that 257 of those neurons are not-Gabor in old age.

| **Gabor-like in youth (30 loops)** |  | **Gabor-like in old age (80 loops)** |
| --- | --- | --- |
| 123/400 |  | 26/400 |
|  | **Gabor-like in old age** | **not-Gabor in old age** |
| **Gabor-like in youth** | 6/123 | 117/123 |
| **not-Gabor in youth** | 20/277 | 257/277 |

### Performance changes

As a first basic check of network performance, we analyze the distribution of spike counts over image presentations. The single change we made to the network—increasing the excitatory target firing rate *p_E_* with training—is intended to produce increased neuron activity with age, as observed in experiments [34]. Indeed, as shown in Fig. 4A, we find that not only do the mean excitatory spike counts increase with age, but the spread of the distribution does as well, confirming that our network exhibits growing excitation during the aging process. We do not increase the target firing rate *p_I_* of the inhibitory neurons, and as a result neither the mean spike counts nor variance of the spike count distribution of these neurons increase with age (Fig. 4B).

**Figure 4:**
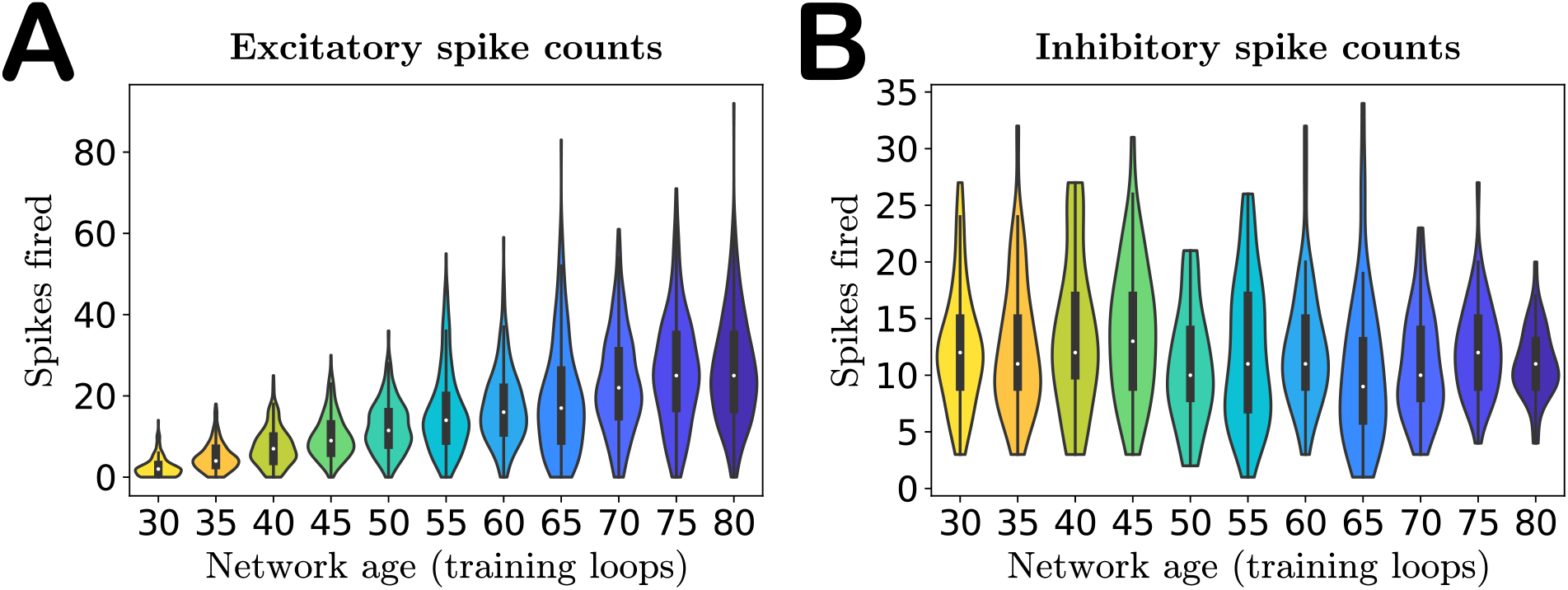
Activity of the network with age. **A**. Violin plot of the excitatory spike count distributions with age. Markers indicate the median. Both median and variance of the distributions increase with age, indicating increased excitation in the network, as expected. **B**. Violin plot of the inhibitory spike count distributions. The inhibitory target spike rate does not increase with age, and thus the distributions do not change systematically during aging. The colors of the violins represent the age indicated on the horizontal axis.

To evaluate functional performance at each age, every 5 training loops we pause training and simulate the network’s responses to patches drawn from pixel images of oriented gratings of spatial frequency 0.1 cycles per pixel, such that one 8 × 8 patch contains approximately 1 cycle. We quantify a neuron’s selectivity to each orientation using the same orientation selectivity index employed by Hua *et al.* [34]. As seen in Fig. 5, the cumulative distribution plots of the orientation indices (the fraction of cells with indices less than a particular value) qualitatively agree with the experimental observations of [34] in young and old cats: selectivity decreases with age, demonstrated by the shift of the cumulative plots toward lower orientation indices. However, our model results also suggest this process may not be a straightforward monotonic degradation of selectivity with age. In the first several epochs of aging our network we find that the mean selectivity of the network does not actually change appreciably, but rather that changes in weakly- and strongly-selective neurons initially balance out (ages 30-50, Fig. 5C), and only later in age does the network begin to lose strongly-selective neurons, a prediction that could be tested experimentally in future studies.

**Figure 5:**
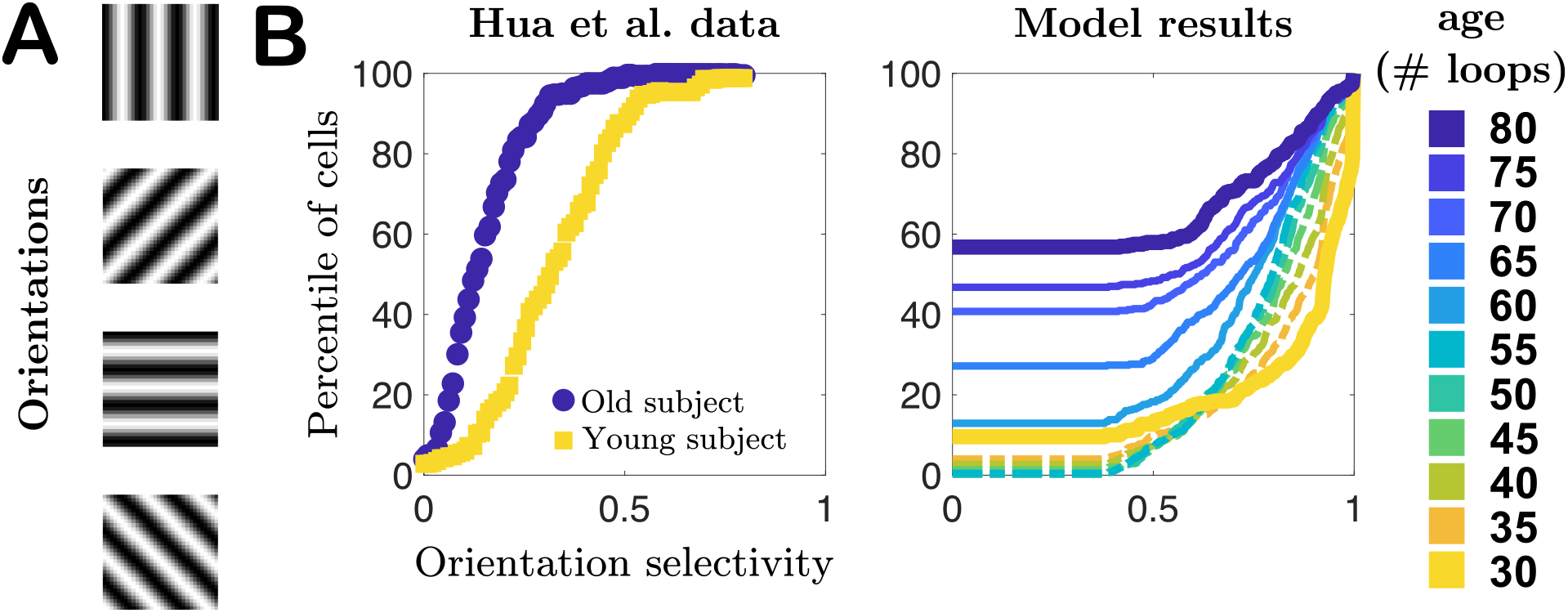
Selectivity of cells to oriented gratings: **A**. Schematic of oriented gratings presented to the network to evaluate selectivity to oriented features in visual input. **B**. Cumulative distribution plots of orientation selectivity indices of real neurons from Hua *et al.* [34] in young (yellow) and elderly (blue) cats, compared to our model results for a series of different “ages” of our network model. In both cases orientation selectivity decreases with age. Additionally, our network model predicts that mean selectivity initially changes little in the early stages of aging.

### Numerical experiments to test contribution of different physiological parameters to declines in performance

Given the variety of changes our model network undergoes during aging, can we identify which physiological changes contribute most to the observed changes in performance? We certainly expect the degradation of the Gabor-like structure of the receptive fields to play a strong role, at least later in the aging process when the RFs are no longer Gabor-like, as in the absence of these edge-detector filters in the network we naturally expect selectivity to drop. However, before the RF structure has completely degraded, we may also expect lateral weights to play a role, as several theoretical and experimental studies suggest that neurons with similar response properties will effectively mutually inhibit one another and thereby sharpen the selectivity of their responses [44–50]. Thus, a loss of inhibition could also contribute significantly to decreased orientation selectivity at intermediate older ages. To probe which parameter set dominates the contribution to the loss of orientation selectivity as a function of age, we performed several numerical experiments on our networks.

In our first set of simulation experiments we age our network while shutting off the learning rules for one or more of the physiological parameters, freezing these parameters to their young values. In this way we can assess how much freezing each parameter set to its young values rescues the orientation selectivity. We perform three such tests: i) we freeze the input weight matrices *Q*, ii) we freeze the lateral weight matrices *W*, and iii) we freeze both *Q* and *W*. Note that we do not freeze the thresholds to their young values because our network aging process is driven by the increasing excitatory target spike rate *p_E_* (*t*_loop_), which only appears in the threshold learning rule. Freezing this learning rule would therefore preclude aging in our network.

The results of these tests appear in Fig. 6, in which we plot the average selectivity as a function of age for each case. We see that both cases in which the input weights *Q* are frozen retain a high mean orientation selectivity across ages, comparable to the mean selectivity in youth. When only the lateral weights are frozen and the input weights are allowed to adapt to the increasing *p_E_* (*t*_loop_), the mean selectivity maintains its youthful value for most of the aging process before suddenly dropping at the oldest ages we simulated, though not quite as low as the case in which both input and lateral weights are reorganized by ongoing plasticity. These results suggest that changes in input weights during aging are more important than changes in lateral weights for maintaining the functional performance of the network; see also our supporting analyses in “Evaluating network selectivity in hybrid networks of mixed young and old parameters.” However, the fact that it is only in advanced age that selectivity falls off when lateral weights are frozen suggests that the coordinated reorganization of both input weights and lateral weights leads to large degradation in functional performance.

**Figure 6:**
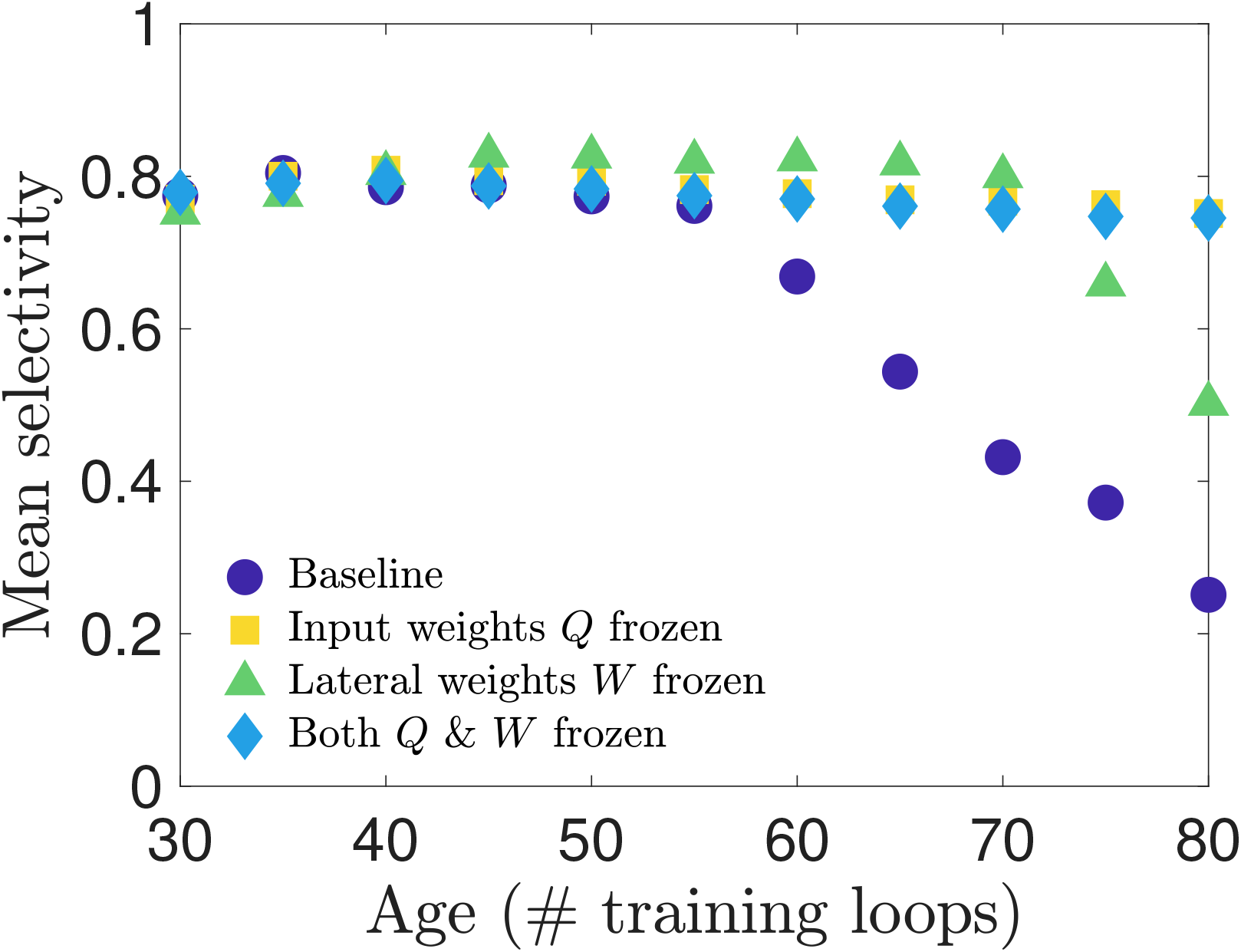
Selectivity of cells when connection weights are frozen during training: We test how network parameters contribute to orientation selectivity by shutting off learning of the input weights or lateral weights during the aging process. When input weights are frozen to their young values, the selectivity is barely impacted. However, when lateral weights are frozen to their young values, the selectivity remains high for most ages until it suddenly begins to drop off near the oldest ages in our simulations. When both Q and W are frozen the selectivity remains near its young values.

These tests so far only demonstrate that input weight changes are the dominant contributor to the decline in selectivity, but do not disambiguate the contribution of changes in input weight magnitude from the contribution of changes in the spatial organization (the receptive field structure). As observed in Figs. 2 and 3, we know that both the receptive field structure and the distribution of input weight magnitudes are altered during our aging process. With our network model we can separate the effects of input weight magnitude versus structure to test whether one of these features contributes more to the changes in neural selectivity. We expect, for instance, that the loss of Gabor receptive field structure will result in diminished orientation selectivity. We test sensitivity to input weight structure and strength in two ways: i) we preserve each individual neuron’s young receptive field structure while remapping the input weight magnitudes such that the distribution matches that of the old-age input weights; ii) we preserve each individual neuron’s aged receptive field structure while remapping the input weight magnitudes to match the young input weight distribution. We find, as seen in Fig. 7, that restoring the young RF structure gives an overall boost to the mean orientation selectivity at all ages, as might have been expected. However, we also see that at the oldest ages (70 to 80 training loops) restoring the young magnitudes yields a greater improvement in selectivity than restoring young structure.

**Figure 7:**
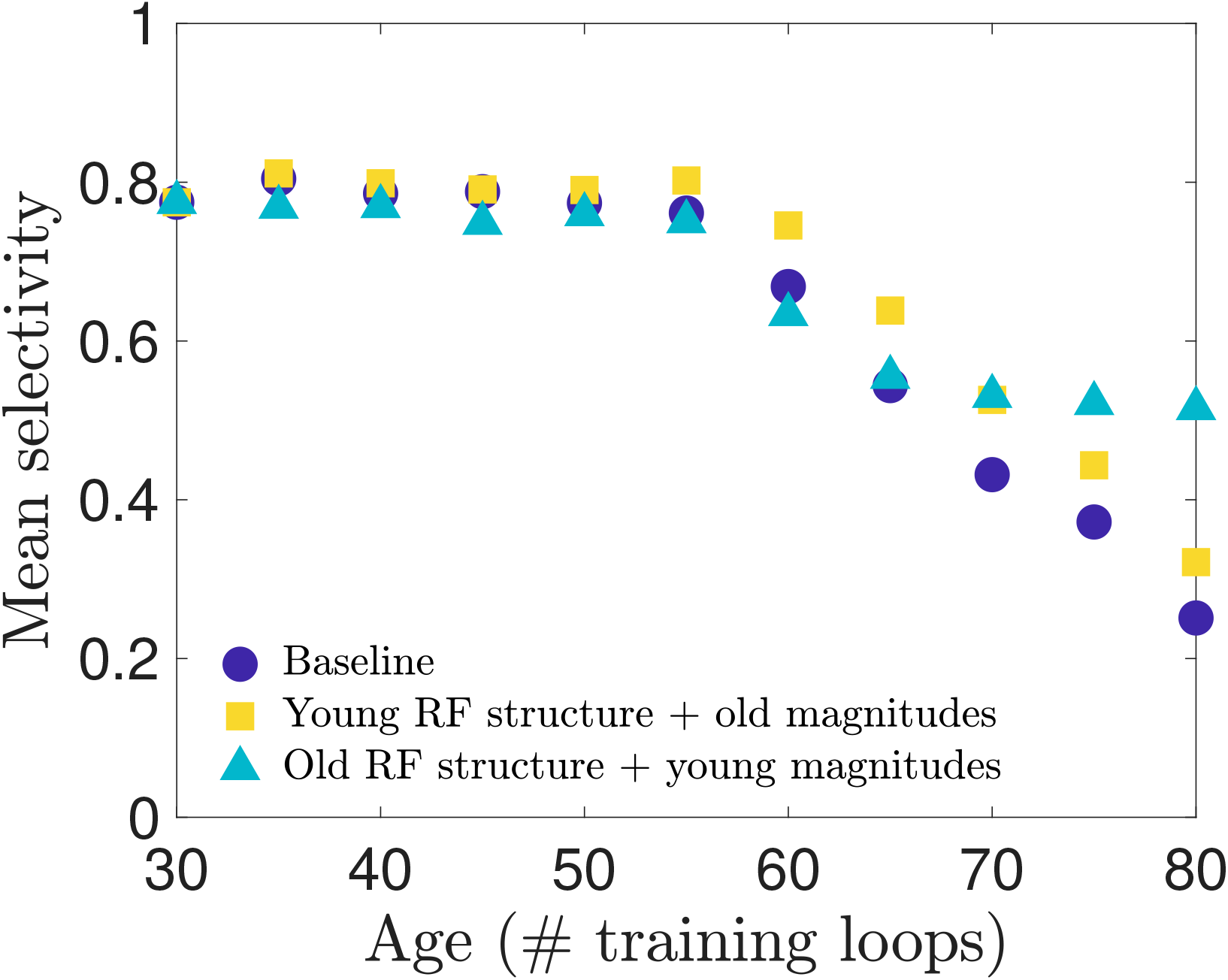
Selectivity of cells with remapped receptive fields: To disambiguate the relative impact that aging receptive field (RF) magnitude versus spatial structure have on orientation selectivity, we remap the magnitudes (input weight values) of each neuron’s RF to create two types of RFs: RFs with the original spatial structure they have in the young network but a magnitude distribution matching the aged distribution, and RFs with the aged spatial structure but young magnitude distribution. (See Methods for details). We create these hybrid RFs for each age and simulate the network model to measure the orientation selectivity of the neuron population. We find that restoring the young RF spatial structure improves selectivity in the early stages of aging, outperforming the baseline network (no remapping) as aging progresses, but the aged RF spatial structure combined with young RF magnitudes maintains an overall higher level of selectivity in the latest stage of aging simulated.

## Discussion

We have demonstrated that if the target spike rate *p_E_*(*t*_loop_) in the learning rule for neural thresholds begins to increase after the network has reached maturity the resulting training process produces networks that qualitatively replicate features observed in senescent brain tissue. The training period of E-I Net’s predecessor, SAILnet [39], has previously been used to model the experiential development of visual cortex in growing ferrets [38], and our work continues this line of thinking by extending the interpretation of the modified training dynamics to aging processes. Though we do not model the underlying mechanism that causes *p_E_*(*t*_loop_) to drift, we interpret this change as arising from a disruption in the homeostatic regulation of this target excitatory spiking rate. Coupled with ongoing synaptic plasticity, this drift of *p_E_*(*t*_loop_) reorganizes the physiological properties of a network in a multitude of ways. The first is an increase in excitation (Fig. 4); this is the intended consequence of altering *p_E_*(*t*_loop_) to increase with continued training. All remaining changes in network physiology, including the weakening of synaptic weights, broadening of firing threshold distribution (Fig. 2), and deformation of the neural receptive fields (Fig. 3) follow from this single modification of the network learning rules. These physiological changes result in declines in functional performance of the network, such as reduced sensitivity to oriented features in visual input (Fig. 5). Moreover, with access to all our model parameters we can show that declines in selectivity are primarily due to the changes in input weight structure and strength (the receptive fields), rather than changes in lateral inhibition (Figs. 6 and 7). These changes are consistent with experimental studies probing aging visual cortex, suggesting that such firing rate dysregulation is a potential mechanism underlying senescence in neural circuitry.

To support this assessment of our results, below we compare our results to current experimental observations in detail and make predictions for future experiments that could lend more evidence toward or against this hypothesized mechanism (Comparison to experimental results and implications for future experiments), and then we discuss the limitations of the model; i.e., which features we expect to be reasonably realistic, albeit simplified, and which features of experimental observations we were not able to replicate with the current model (Limitations of the model), before giving a final summary and conclusion (Summary of results and implications for disease).

### Comparison to experimental results and implications for future experiments

Our investigation was originally motivated by several experimental results in visual cortex: i) spontaneous and stimulus-evoked neural activity increases with age [34], ii) the proportion of GABAergic neurons in senescent brain tissue decreases with age (but total neuron density does not) [35, 36], and iii) aged neurons exhibit decreased orientation and direction selectivity.

It is not clear *a priori* what the causal relationships are between these observations; most likely, there is an underlying common cause, rather than several (approximately) independent causes leading to these changes. Intuitively, we might expect that decreases in GABAergic inhibition could lead to increased firing in the network. We explored this hypothesis in an early version of this work by training the network model to maturity and then turning off learning and decreasing the magnitude of inhibitory connection weights in the network by a global fraction. We were able to reproduce loss in performance only when we added a tonic current to drive the neurons, and only for particular ranges of this tonic current and global reduction in the inhibitory weights. However, as the changes to the inhibitory weights in this early version of the model were implemented by hand, and not by changing the learning rules, they provided no plausible mechanisms for explaining *how* the network aged. Moreover, because learning was turned off in this approach, it did not allow for the possibility that other network parameters might adjust to compensate.

We ultimately developed the hypothesis pursued in this work, that the causal relationship could be that a breakdown in homeostatic regulation of target excitatory firing, coupled with long-term synaptic plasticity, would lead to changes in synaptic strength and other physiological features, as well as changes in functional performance. Therefore, the fundamental change we made to the healthy mature network was intended to produce increased activity by steadily bringing up the excitatory target spike rate as training progressed past the phase at which the network parameters had reached a steady state. All other results follow from the coupling between this change and the learning dynamics of the model parameters. As expected, the mean firing rate of excitatory neurons increased with network age, as did variance of firing, as shown in Fig. 4.

The consistency of the observed decreases in synaptic weight magnitudes in our model (Fig. 2) with experimental observations in visual areas is more subtle and open to interpretation. The direct experimental observation in [35,36] is that the density of GABAergic neurons measured by GABA-immunoreactive staining decreased in older subjects compared to younger subjects; however, the total density of neurons (measured by Nissl staining) *did not* decrease. This interpretation is supported by other work [51], which finds minimal neuron loss with age and notes that during aging genes involved with GABA-mediated neurotransmission are strongly down-regulated in human and macaque pre-frontal cortex. Thus, observed age-related changes are not caused by a loss of GABAergic neurons through cell death, but more likely a drop in GABA expression itself. In a conductance-based leaky-integrate-and-fire model this could be interpreted as a decrease in the maximum fraction of GABAergic receptors being activated by a pre-synaptic spike; as E-I Net is a current-based leaky-integrate-and-fire model, the closest equivalent effect is a change in synaptic weight. Because the reversal potential of GABAergic synapses is typically much larger in magnitude than the membrane potential, it is often valid to approximate the inhibitory conductance-based synapses as current-based synapses [52]. Therefore, we interpret the observed decrease in the range of inhibitory input and lateral synaptic weights in our model to be consistent with the experimental observations.

In addition to the observed decreases in GABAergic neurons [35], experimental work studying postmortem tissue from human visual cortex [8] has also found evidence that the density of several types of glutamatergic receptors tends to decrease in old age, a trend which also appears to hold across several species and brain areas, with some exceptions [5,6]. We interpret these decreases in glutamate concentration and/or receptor density as being consistent with the observed decrease in excitatory synaptic weights in our model (Fig. 2). We were not aware of these results during the original formulation of our model, so the qualitative agreement between these studies and our model’s decrease in excitatory weights in addition to inhibitory weights may be viewed as a post-diction of our model.

As with the inhibitory synapses, there does not exist a perfect translation of experimental observations on excitatory neurotransmission into expected changes in our current-based leaky-integrate-and-fire model, but it is reasonable to interpret a decrease in receptor count or neurotransmitter concentration as resulting in an effective decrease in synaptic conductances, which approximately map onto the synaptic weights Wij of our model when neglecting the voltage dependence of the conductance. Some additional caveats apply to this approximate mapping for excitatory neurons compared to inhibitory neurons, as there are several principal excitatory neurotransmitters to consider, each of which could change in different ways with age. The most consistent finding in [5] was a decrease in the density of NMDA receptors with age across several brain regions. Another possible caveat in our conductance-to-current mapping is the fact that the true conductance-based nature of excitatory synapses could become relevant because excitatory reversal potentials are not significantly larger than membrane potentials, so a current-based approximation would not fully capture changes in excitatory synapses with age if they were to cause membrane potentials to cross the excitatory synaptic reversal potentials. However, to the extent that a current-based integrate-and-fire model can qualitatively capture these changes, we believe it is reasonable to interpret our results as replications of these experimental findings.

In addition to the weakening of synaptic connection weights, our model also predicts a change in the firing thresholds of neurons. Because E-I Net uses non-dimensionalized membrane potentials, changes in model thresholds could correspond to changes in the physiological thresholds *or* resting potentials. Data on how firing thresholds and resting potentials change with age do not appear to show any consistent trends across species or brain areas, with several studies finding no apparent changes in many cases [7, 53], some finding increases in firing threshold [7], while others find increased hyperpolarization of resting potentials and firing thresholds [54]. Taken at face value, the increase in the spread of firing thresholds in our model (Fig. 2) suggests that even within a brain area there may not be a consistent trend, though further study is needed to test this possibility.

In terms of the loss of functional performance in older brain tissue, our aged network model qualitatively replicates Ref. [34]‘s observed loss of orientation selectivity to grated stimuli compared to the mature young network. While there are quantitative differences between our model predictions and the experimental observations of [34], these quantitative differences between our simulation results and the data may be traced to some of the simplified features of the network model we use. For example, the selectivities of neurons in our model network are much more polarized, having many neurons with vanishing orientation selectivity index (as they did not fire to any stimulus when presented with patches from the oriented grating images) and many with nearly maximal selectivity, which also leads to our cumulative plots having the opposite convexity to that of the experimental data. These more polarized orientation selectivities in our simulated network are produced in part because the learning rules of E-I Net promote sparse firing yet demand decodability of inputs [39], resulting in highly selective neurons that are active essentially only when the feature they are selective for is presented, and are silent otherwise. Moreover, because the network is deterministic (there are no noisy current inputs), the resulting activity levels tend to be highly polarized. Intermediate selectivities are likely generated by lateral coupling between neurons with similar but distinct receptive fields [42], leading to imperfect selectivity. Another factor that promotes these more polarized selectivities is the fact that every neuron in the network responds to the same visual patch at a given time. Potentially, a very large network in which different clusters of neurons respond to many inputs from different locations in the visual field would generate enough recurrent activity for recorded neurons to appear spontaneously active, which might smooth out the distributions of orientation selectivity. We leave training and simulating aging in such a network for future work.

As our model gives us full access to network parameters, we were able to demonstrate that this decline in orientation selectivity is primarily caused by the deterioration of Gabor-like receptive fields of the model neurons as the runaway target excitation causes the network to reorganize, rather than, say, by changes in lateral weights that might also have been predicted to impact sensitivity [44–50]. In particular, we found both that the net proportion of Gabor-like receptive fields decreased as the network aged (Table 1) and that the receptive field of a given neuron was not preserved with age (Fig. 3). The net loss of Gabor-like receptive fields impacting orientation selectivity makes intuitive sense, as such receptive fields are typically interpreted to be edge-detectors [55]. This is also a new prediction that could potentially be tested experimentally in longitudinal experiments on the same individuals, measuring the receptive fields of populations of neurons in visual cortex in both youth and old age, inferring the receptive fields using, for example, generalized linear model fits [56–61], and comparing the distributions of Gabor-like versus not-Gabor receptive fields at these different ages.

Other experimental work has found that contrast sensitivity is diminished with age [2,62]. We did not perform a detailed investigation of whether our model could qualitatively replicate this result, for technical reasons: images on which the network model is trained are first “whitened” to remove the mean intensity of the images and normalize the variance across pixels in an image—operations interpreted within the context of E-I Net to be performed by the retina and lateral geniculate nucleus (LGN) en route to V1. However, this pre-processing step will also normalize the contrast of an image. To properly test contrast sensitivity would thus require a model of the aging retina and LGN, which is beyond the scope of this work, but could potentially be developed and implemented as a modification of the whitening step of this model, or perhaps studied with a similar setup, using recent deep network models of retinal circuitry [63, 64]. In addition to retinal changes, we have not attempted to account for the variety of potential age-related physical changes in the lens or vitreous fluid of the eye, which can also alter the image information transmitted to visual cortex. However, modeling how visual input transforms with age-related changes in the eye [65] may allow future work to better estimate how these effects might influence the aging process in models like ours.

### Limitations of the model

Although our model successfully replicated several experimental observations and makes predictions for new experiments, there are of course some results the model as currently implemented could not replicate. Moreover, as a simplified model of network development and dynamics, there are features of the model that could be considered biologically implausible or oversimplified, and warrant discussion here.

One of Ref. [34]‘s results that our model did not successfully reproduce is the loss of direction selectivity. We trained our network on sequential patches of image frames from the CatCam database [66, 67] for the purpose of evaluating direction selectivity, but found that our network was not complex enough to learn direction selectivity from training on these natural movies alone. Our network was able to learn some direction selectivity when trained on moving grating stimuli, but this began to overwrite the network structure learned from the natural scenes. Because we interpret the trained network model as a computational stand-in for mammalian V1 that developed in natural environments, we considered it invalid to induce direction selectivity by training on moving gratings before probing network responses.

The difficulty of learning direction selectivity could be because directed motion is not represented frequently enough in our training movies, but could also be because our network neurons exponentially filter inputs, all with the same single timescale. Potentially, a modified network model that allowed for different spatiotemporal history filters or even multiple membrane timescales might be better suited to learn direction selectivity from natural movies. A model that could successfully learn direction selectivity may also prove useful for testing how age impacts neural decoding of stimulus speed, which has also been shown to deteriorate with age [68].

The biological plausibility of training neural networks has been debated extensively in the literature [69, 70], casting doubt on whether training such networks can be an appropriate model for network development. Many, if not most, of these criticisms focus on training algorithms that employ non-local learning rules (i.e., rules which require global information about the network in order to update an individual neuron’s synaptic connections), such as backpropagation [71]. However, the learning rules employed by E-I Net and its predecessor SAILnet [37,39] are local (Eqs. (2)–(4)), meaning that updating a particular neuron’s parameters only requires information about its own incoming connections and firing threshold. Therefore, the learning rules used in our model may be considered a reasonable simplification of biologically plausible learning mechanisms, and thus can capture the essential minimal features of network development. However, it is worth noting that the learning rules of E-I net do not give rise to any layered or modular network organization, so the resulting network is best interpreted as a particular layer of V1, such as layer 4, rather than an entire cortical column.

One subtle and potentially unintuitive feature of our model is the fact that the time-course of training need not be straightforwardly mapped onto the time-course of development. For instance, we use the same learning rates for the entire duration of our training procedure, including both the development phase (loops 1 – 30) and our aging phase (loops 30 – 80). However, experimental evidence indicates that there is a “critical period” in which long-term synaptic plasticity is high early in development, after which plasticity drops, allowing the nervous system to approximately lock in learned structure and prevent sudden drastic changes in sensory experience from overwriting this structure [72]. At first, a critical learning period seems at odds with our training procedure, which has no such period, but that is true only if one assumes a linear relationship between training time and true organism age. A nonlinear relationship would allow for our results to be consistent with a critical period. In particular, we could assume that training time and organism age are approximately linearly related during development (loops 1 — 30) but undergo a shift to a different linear relationship after maturation that holds during the aging phase. We have checked that decreasing all learning rates by a factor of 10 and extending stimulus presentation time by a factor of 10 after 30 training loops indeed give results qualitatively similar to those of the baseline model; see Critical learning periods and time rescaling in Methods).

Finally, our results do not rule out other possible mechanisms for generating the observed changes in networks with age, nor do they explain what physiological mechanisms might cause a drift in target firing rate to begin with. Possible causes could be related, for example, to changes in calcium dynamics controlling synaptic plasticity [73], but a detailed study of such possible breakdowns of this firing rate regulation are beyond the scope of the current work. Nevertheless, starting from the hypothesis that there is some breakdown of homeostatic regulation of target firing rates, causing them to increase with age, in the presence of ongoing long-term synaptic plasticity has allowed us to replicate several experimental results and make predictions for effects not yet observed experimentally. Future experiments testing these predictions would enable validation or refutation of this hypothetical mechanism.

### Summary of results and implications for disease

We have shown that altering the target firing rate of neurons in a trained V1 network model—interpreted here as the result of a dysregulation of homeostatic mechanisms regulating network firing rates—amid ongoing long-term synaptic plasticity results in several physiological and functional changes symptomatic of aging in visual cortex. Our single modification to the learning rules of the E-I Net model of visual cortex is intended to promote increased firing with age in the network, and successfully reproduces this feature. The other features replicated or predicted by our approach follow from this single modification. The replicated features include

- reduced synaptic strengths, in the form of weakened input connections to the excitatory and inhibitory subpopulations as well as diminished lateral connections between and within these subpopulations, and
- impaired stimulus-feature detection, in the form of decreased selectivity for oriented grating stimuli, while predicted features include
- a net loss of Gabor-like receptive field structure of V1 neurons, and
- the principal role of a decline in strength (i.e. input weight magnitude) and Gabor character of receptive fields in the loss of orientation selectivity; decreases in lateral weight strength also contribute, but not as significantly.

Thus, our work suggests that dysregulation of network excitation may be one of the causal mechanisms that leads to physiological and functional changes in networks and their dynamics as an organism ages. While this insight does not rule out other possible mechanisms, it provides a basis for forming new hypotheses and devising experiments to test these ideas, either by looking for experimental evidence of model predictions in naturally aged neural tissue or perhaps by testing whether it is possible to induce senescent neural phenotypes in visual cortex by disrupting firing rate homeostasis. Future studies could test this mechanism either by performing new recordings in aged tissue to compare to our model predictions, or by direct experiments in which firing rate homeostasis in young tissue is actively disrupted by the experimenter to assess whether some features of old-age phenotypes develop. Should further studies provide evidence that this mechanism contributes to aging processes in *vivo,* our results also suggest possible ways some of these performance changes could be partly mitigated. For instance, our numerical experiment varying the structure and strength of input into visual cortex (Fig. 7) suggests that improving orientation selectivity in aging patients could potentially be accomplished by coarsely increasing the gain of relevant clusters of neurons. In advanced age this approach could potentially be more effective than targeted treatments that would aim to precisely restore the inputs to individual neurons, a more complex and potentially invasive treatment.

Beyond replicating previous experimental observations and making predictions for new experiments, modeling how neural circuitry changes with age will play an important role in understanding the progression of neurological disorders whose onset correlates with age, such as Alzheimer’s disease or Parkinson’s disease, as well as general age-related cognitive decline. In particular, having a baseline model for predicting changes in healthy neural circuitry is important for disambiguating which effects are symptomatic of disease from those likely to be present in a healthy aging individual, and thereby determining which aspects of the disease progression to focus on and develop treatments for.

## Methods

### Network model

We use E-I Net, a spiking network model of V1 trained on natural images, as the basis for our investigations. E-I Net is written in MATLAB, and all modifications and subsequent analysis code are also in MATLAB. We will give a brief overview of the model here, and then focus on modifications of the model we implemented for use in this study. Some of these modifications were non-default options already available in the code, while others are modifications we developed ourselves. For full details on E-I Net we direct readers to [37].

E-I Net is a network of leaky integrate-and-fire neurons that spike in response to pixel-image inputs *X_k_*. In discrete time the membrane dynamics of the network are given by

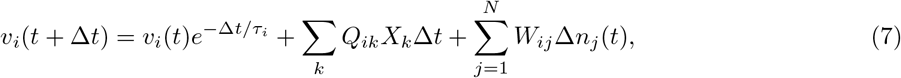

with spiking conditions

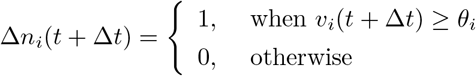

followed by a reset

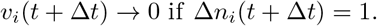

These equations are the discrete-time version of Eq. (1) used in practice by [37]. Here, Δ*t* is the time-step size, *τ_i_* = *τ_E_* or *τ_I_* is the membrane time constant of neuron *i,* and Δ*n_i_*(*t*) = *n_i_*(*t*) – *n_i_*(*t* – Δ*t*) is the number of spikes fired by neuron i on time-step *t*. Neuron *i* fires a spike on time-step *t* if *v_i_*(*t*) ≥ *θ_i_*, after which its membrane potential is reset to *v_i_*(*t* + Δ*t*) = 0. The sum indexed by *k* is over the pixels of the input patch *X*, and the sum *i* = 1 to *N* is over the *N* neurons of the network. The network is partitioned into excitatory and inhibitory subpopulations; for our simulations reported in this work, we follow [37] in using *N_E_* = 400 excitatory neurons and *N_I_* = 49 inhibitory neurons. In this paper we take *W_ij_* to be positive (negative) if the post-synaptic neuron *j* is excitatory (inhibitory), in contrast to the original E-I Net paper, which makes the sign explicit, such that *W_ij_* is non-negative. We do not include excitatory-excitatory lateral connections (i.e., *W_EE_* = 0), as we expect such connections to be negligible physiologically; see [37] for additional details and justification of this choice and other aspects of the training of the model.

The network learns input weights *Q_ik_*, synaptic weights *W_ij_*, and firing thresholds *θ_i_* according to the learning rules given in the main text (Eqs. (2)-(4)). For the reader’s convenience, we repeat these learning rules here:

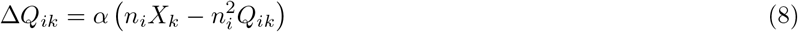

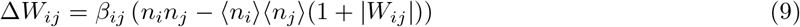

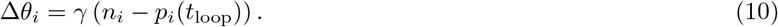

These learning rules impose linear decoding of the image inputs (Eq. 8), minimize redundancy in spiking (Eq. 9), and promote sparsity in firing (Eq. 10). As in the main text, *n_i_* is the spike count of neuron *i* in response to the presented image patch over a duration *T*, (*n_i_*) is a moving average over time, weighted by an exponential decay factor, and the parameters *α*, *β_ij_* ∈ {*β_EI_*,*β_IE_*,*β_II_*}, and *γ* are the learning rates. The target number of spikes per time-step is *p_i_*(*t*), equal to a constant *p_I_* for inhibitory neurons and an “age”-dependent function *p_E_*(*t*_loop_) for excitatory neurons, where

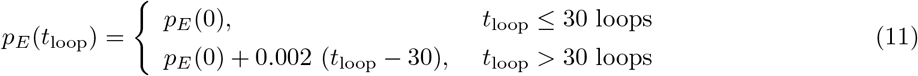

as in the main text. The “time” *t*_loop_ corresponds to the number of training loops that have taken place (the network’s “age”). The moving averages involved in the computation of 〈*n_i_*〉 are over the time-course of the network simulations at each age.

The standard values used for the parameters in our network training and simulations are given in Table 2. For the learning rates we follow the original E-I Net model for our baseline cases. These rates are modified in the numerical experiments discussed in the section “Numerical experiments to test contribution of different physiological parameters to declines in performance” by setting one or more of these rates to 0. In the E-I Net code, these rates are combinations of several parameters that we do not list here; however, we note that the numerical values we give include a global learning factor, a multiplier applied to all the learning rates, which is set to 0.4, the value used for accelerated training shown in examples in the E-I Net code repository [37]. We summarize all of the parameters used in the model in Table 2.

**Table 2:** Parameters and standard values used in this work: This table summarizes the parameters that appear in the equations we give in the main text, along with the standard values we use in our baseline simulations.

| Parameter | symbol | value |
| --- | --- | --- |
| Number of excitatory neurons | $N_E$ | 400 |
| Number of inhibitory neurons | $N_I$ | 49 |
| Input weight learning rate | $\alpha$ | 0.008 |
| $E \leftarrow I$ lateral weight learning rate | $\beta_{EI}$ | 0.028 |
| $I \leftarrow E$ lateral weight learning rate | $\beta_{IE}$ | 0.028 |
| $I \leftarrow I$ lateral weight learning rate | $\beta_{II}$ | 0.06 |
| threshold learning rate | $\gamma$ | 0.028 |
| Initial excitatory target spike count | $p_E(0)$ | 0.01 |
| Inhibitory target spike count | $p_I$ | 0.05 |
| Network simulation duration (# time-steps) | $T$ | 50 |
| Excitatory membrane time constant | $\tau_E$ | 0.9491 |
| Inhibitory membrane time constant | $\tau_I$ | 0.9491 |
| Time-step size | $\Delta t$ | 0.1 |

The network learning rules are entirely local [37,39] (i.e., changes to a neuron’s parameters depend only on inputs that neuron receives), and hence admit a straightforward biological interpretation as plastic or homeostatic mechanisms that shape network wiring or firing. When trained on natural image inputs the network has been shown to learn realistic Gabor-like receptive fields and produce activity that can be used to accurately reconstruct input [37, 39].

### Training the network

Training our network model consists of two phases: the initial development phase, during which the network forms, and the aging phase during which we implement modifications to the learning rules to cause changes in network structure that mimic the effects of aging. The first phase of training is largely the same as the original E-I Net training procedure, while the aging phase is a novel modification that we implement after the first phase reaches a steady state.

SAILnet [39] and E-I Net [37] train their networks on static pixel images of natural scenes from the database of images used by Olshausen and Field in their seminal work on the emergence of Gabor-like receptive fields under sparse-coding principles [74]. Patches of these scenes are chosen randomly from a given movie in this database and each patch is presented to the network for 50 time-steps of simulation of the leaky integrate-and-fire network model. Following each image presentation, the state of the network’s membrane potentials are reset. To speed up training, E-I Net creates 100 copies of the network, presenting each copy with a different set of randomly selected patches from the movie. At the end of the training epoch, the spiking data from all 100 copies is amalgamated to determine how the network parameters should be updated.

We modify this procedure slightly in training our networks. We train our networks on natural scenes from movies in the CatCam database, which depict the scenes encountered by a cat exploring a forest [66, 67]. We choose the CatCam database because our original intent was to investigate changes in both direction selectivity and orientation selectivity, as noted in the Discussion. For this purpose, it is necessary to train the network on temporal *sequences* of patches. We use 50 contiguous frames of 8 × 8 patches from these natural movies. We present each patch in this sequence for 50 time-steps of network simulation before presenting the next patch in the sequence. We do not reset the membrane potentials of the neurons between patches (an option available in E-I Net). We similarly use movies of grating images to evaluate the orientation selectivity of the network, taking 40 frames of gratings drifting perpendicularly to their orientation, covering one direction for each of the four different orientations; we ceased evaluating performance on both grating directions after finding that the networks did not develop significant direction selectivity. Although the networks we report on in this work do not ultimately develop significant direction selectivity when trained on natural movies, we keep the sequential presentation procedure for training our networks.

In our model we take the fully trained network to represent our mature “young” network, trained for 30 loops at the baseline excitatory target spike rate of *p_E_* = 0.01 expected spikes per image presentation and *p_i_* = 0.05 expected spikes per image presentation for inhibitory neurons, comparable to the values used in [37], which use *p_E_* = *p_I_* = 0.05 for the entire duration of training. The goal of this work is to modify this model to reflect experimentally observed age-related physiological changes, such as increased neural activity and decreased inhibition, and thereby produce “aged” networks in which we can test how functional performance and coding have changed. Rather than implement the physiological changes by several independent by-hand adjustments, a much more parsimonious modification is to alter only the excitatory target spiking rate *p_E_*. If we increase *p_E_* with training time past the mature young stage of the network, the network will reorganize to achieve a higher mean firing rate close to this new target. We increase *p_E_* by 0.002 every subsequent loop, culminating in the “old” network after 80 total loops. This not only results in increased excitatory neuron activity (both in terms of mean firing and variance of firing), but also naturally gives rise to a decrease in inhibition (Fig. 2). The changes in selectivity to oriented grating stimuli also emerge from this single by-hand change. To check how sensitive the aging procedure is to the initial value of *p_E_*, we also ran two additional network simulations starting from higher values, *p_E_*(0) = 0.05 and *p_E_*(0) = 0.07 spikes per image presentation; see Supplementary Figure 11. Because our method produces a sequence of aged networks that are gradual evolutions of earlier “ages,” we loosely interpret the training process as the “aging” process in our network. (As noted in the Discussion, there is no definite relationship between training loops and organism age). This is similar in spirit to previous work interpreting the initial training phase of the network as a model for the developmental period of visual cortex [38].

### Convergence to steady-state

Our aging process does not begin until 30 training loops have occurred, at which point the network has reached a steady state at which continued training only causes the network parameters to fluctuate around a mean value. In practice, we determine whether the network has reached a steady state using the mean orientation selectivity of the network. We find that by approximately 20 loops the mean orientation selectivity of the network has converged to 0.77 and continues to fluctuate around this value. Thus, the additional 10 loops of training serve as a buffer to ensure the network has reached its mature steady state.

As explained above, after the network reaches this steady state we begin to increase the excitatory target spike rate *p_E_*(*t_rmloop_*) every loop, pausing training every 5 loops only to evaluate the current network’s orientation selectivity. One may therefore wonder whether the network is able to achieve a new steady state during each of these loops (though this is not a major concern, as it is entirely plausible that the aging effects induced by increasing *p_E_*(*t_rmloop_*)) outpace the ability of the synaptic plasticity to keep up. To determine whether a single training loop at the increased value of *p_E_*(*t*_loop_) is enough for the network to remain close to a new steady state, we perform an additional test in which we use a modified procedure in which we increase *p_E_*(*t*_loop_) every loop as in our regular procedure, but then at every 5^th^ loop we hold *p_E_*(*t*_rmloop_) fixed for 5 subsequent loops. We find that these additional loops do not appreciably affect the course of degradation of the network, and network selectivity does not recover. This suggests that, at least after the initial maturation period, the network performance is approximately a state function of the target excitatory spike rate.

### Evaluating network responses to oriented gratings

Every 5 loops we pause training and evaluate the network’s responses to moving grating stimuli (see Fig. 5) of 4 possible orientations in order to measure the selectivity of model neurons, during which we set the global network learning rate to zero so that network parameters do not adapt to the statistics of the gratings. We generate the gratings at spatial and temporal frequencies of 0.1 cycles/pixel and 0.1 cycles/frame, respectively, using the `GratingStim` module of `VisualStimulusToolbox` [75]. As in the training phase, we use 8 × 8 pixel patches of these grating images as visual input to our network. For this patch size there is approximately one cycle of the grating per patch.

For each sequence of grating presentations we record the spiking activity of all neurons in response to grating patches of each orientation and use these responses to compute the orientation selectivity of each neuron. For each of the four orientations, we draw 100 patches from 10 frames of the grating movie. While there are various methods of determining orientation selectivity [76, 77], we choose to follow [34] and quantify the overall selectivity of each neuron to grating orientation as

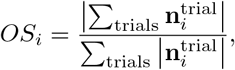

where 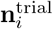 is a vector of the number of spikes neuron *i* emitted in response to each orientation on a given trial (frame presentation). Each trial lasts 200 time-steps of the leaky-integrate-and-fire network dynamics (compared to 50 time-steps during training on the natural movies).

### “Gabor-ness” analysis

The receptive fields (RFs) of the neurons in the network are observed to evolve over the aging process (Fig. 3). Here we define a neuron’s RF as the pixel-space arrangement of the weights *Q* on its connections from pixel-input patches (Fig. 1C). We quantify change in RFs by fitting a Gabor wavelet profile *G* to each RF in both youth and old age. RFs with fits for which the normalized goodness-of-fit, ||*G* – *RF*||^2^/||*RF*||, is less than 0.8 are classified as Gabor-like, while RFs with fits that exceed this threshold are classified as “not-Gabor.” This is the same procedure as performed in [39], except that we use a less strict classification threshold (0.8 vs. 0.5). The Gabor profile takes the following form:

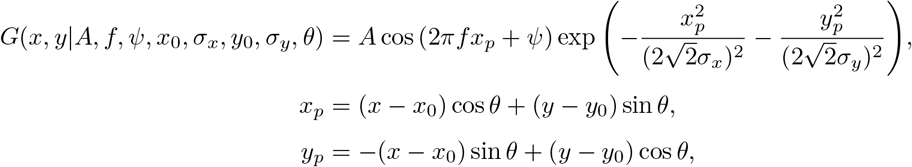

where *A* is the amplitude of the Gabor wavelet, *f* is the frequency of amplitude oscillation, *ψ* is the phase, and *σ_x_* and *σ_y_* are the spreads of the Gabor profile in the *x* and *y* directions. The Gabor wavelet is taken to be a function of independent coordinates *x_p_* and *y_p_*, related to the coordinates *x* and *y* of the pixel-grid representation of *Q* shown in Fig. 1C by an angle of rotation *θ* (not related to the firing thresholds) and offsets *x*_0_ and *y*_0_ corresponding to the shifted center of the Gabor profile. In addition to classifying fits that fail our goodness-of-fit criterion as not-Gabor, we also assign this classification to fits that are centered outside the frame of the receptive field. In principle, we would wholly exclude from the statistics any receptive fields for which fits cannot be computed, but this was never the case for the neurons in our model (see Table 1).

### Numerical experiments

#### Learning rate freezing

To examine the contribution of each network property to the observed decline in neural orientation selectivity as a result of increase in *p_E_*, we conduct the following series of tests. We train several instances of the network to maturity (30 loops) under normal conditions. Upon the completion of the 30^th^ loop, we set the learning rate to zero for different sets of connection weights: in one instance, we zero out the learning rates for only the input connections to the excitatory and inhibitory neurons; in another, we zero out the learning rates for only the lateral connections between neurons; and in the last instance, we zero out the learning rates for both input weights and lateral weights. There is no case in which we zero out the learning rates of the thresholds, as it is through this learning rule that our network ages. After zeroing out the learning rates, we continue training these variants until old age (80 loops), probing selectivity every five loops; see Fig. 6.

#### Receptive field remapping

To decouple the effects of the changing RF structure and magnitude on network selectivity, we examine, independent of the training procedure, the selectivity after remapping the young network’s input weights onto a later distribution of input weight magnitudes, for the networks after 35,40,45...80 loops:

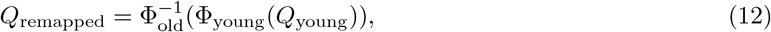

where Φ_young_(·) and Φ_old_(·) are the cumulative distribution functions (CDFs) for the young and old input weight distributions. Eq. (12) modifies the value of every input weight such that the distribution of remapped young input weight *values* matches the distribution of old input weight values, but the spatial pixel organization of each RF is not altered. We empirically estimate the CDFs and inverse CDFs from the data and use interpolating splines to obtain smooth estimates. Thus, we can insert these remapped weights into our old networks to test how RF structure contributes to selectivity. We find that networks with these remapped input weights exhibit consistently higher selectivity than their counterparts with the original aged input weights. We can also test the inverted remapping, 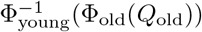, in which the aged networks’ input weights are remapped onto the young distribution of magnitudes, retaining the old spatial pixel structure but with young magnitudes. Networks with these remapped weights are comparable in selectivity to their corresponding original (baseline) networks in the early stages of the aging process, and increasingly more selective late in aging. See Fig. 7.

### Supporting analyses

#### Evaluating network selectivity in hybrid networks of mixed young and old parameters

To support the numerical experiments discussed in “Numerical experiments to test contribution of different physiological parameters to declines in performance,” we also perform a set of experiments in which we evaluate the orientation selectivity of several young-old hybrid networks that we create by mixing together different combinations of the young (age 30) and oldest (age 80) learned parameters. As shown in Fig. 8, the mean selectivity appears to be impacted most when input weights are old and thresholds are young (bars labeled “OYY” and “OOY”). This is most likely because the input weights are smaller in magnitude in old-age than in youth, and therefore the current inputs to each network version ∑_*k*_ *Q_ik_X_k_* are comparatively smaller, whereas the young threshold values are adapted to correspondingly higher current inputs and are therefore typically set higher in order to achieve the target spike rates during development (*p_E_*(*t*_loop_) = 0.01). Thus, typical current inputs drive the neurons to fire less. We see that replacing the young thresholds with the old-age thresholds improves the mean selectivity back to the pure old-age values (bars “OYO” and “OOO”). These results also demonstrate that replacing the young lateral weights with old lateral weights does not appear to have a substantial impact on the mean selectivities at all, as simulations differing only in young/old lateral weights have nearly identical selectivities. Taken all together, this analysis supports the conclusions of our parameter freezing experiments (Fig. 6) that the input weights are by far the most influential contributors to the decline in orientation selectivity, and any sharpening of selectivity that lateral weights might contribute is a second-order effect.

**Figure 8:**
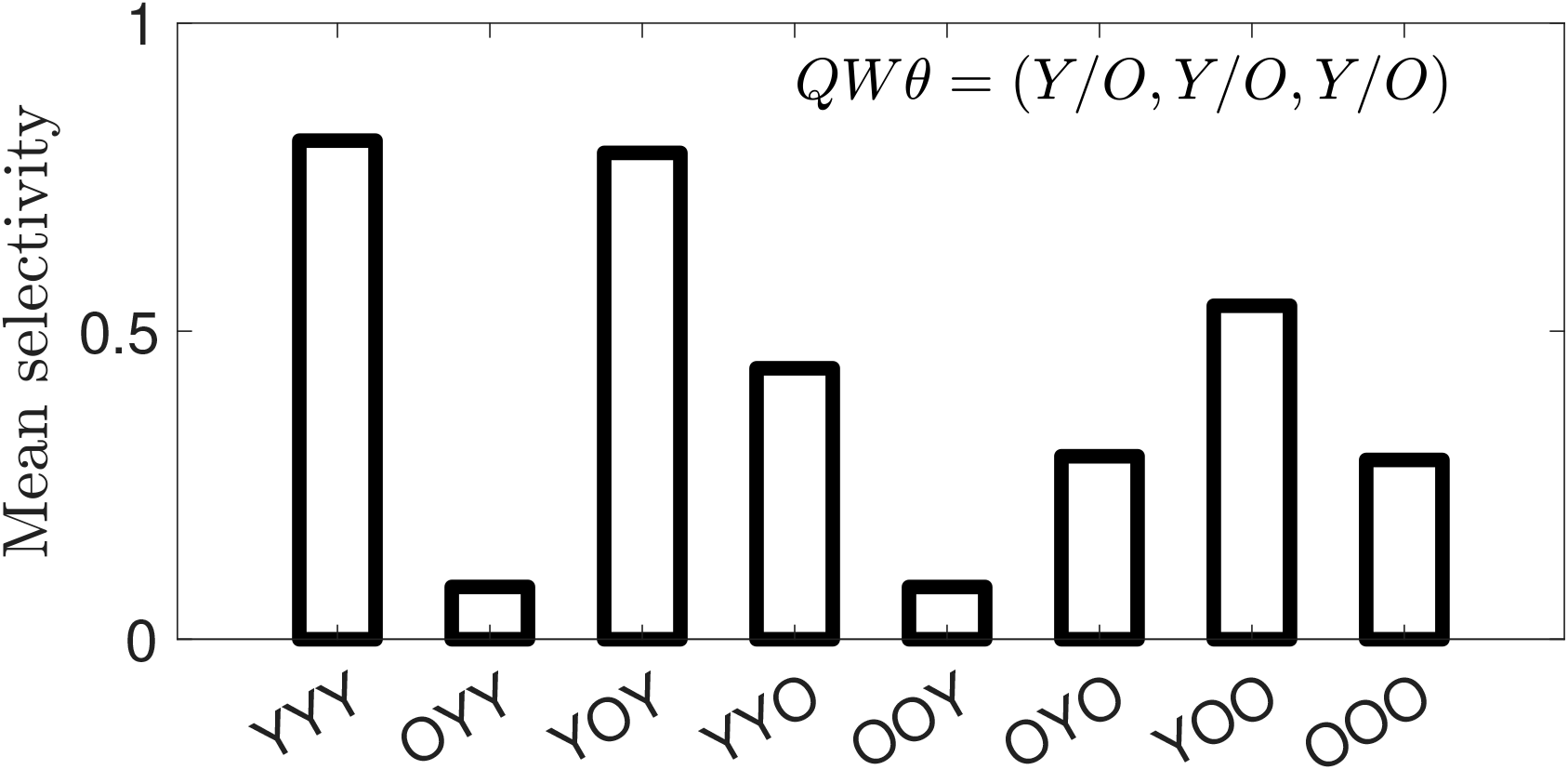
Mean selectivity in networks with swapped parameter sets: The mean orientation selectivity of neurons in network simulations in which we mix-and-match the parameter sets of young (30 loops) and old (80 loops) networks. Horizontal axis labels correspond to young (Y) or old (O) input weights (Q), lateral weights (*W*), and thresholds (*θ*). For example, the “OOY” result corresponds to a simulation using the old-age input weights and lateral weights but the young thresholds.

#### Critical learning periods and time rescaling

The critical learning period hypothesis posits that there is a window early in development in which learning rates are high, which then taper off to lower levels of plasticity [72]. This is unlike our network model, in which the learning rates are the same throughout the training procedure (lifespan). To demonstrate that our training procedure does not preclude interpretation in terms of a critical learning period, we run a training case in which, after 30 loops, we reduce the global learning rate by a factor of 10 (thus reducing all network learning rates by the same factor) and extend the length of natural image stimulus presentation by a factor of 10, and then train the network for 50 more loops under these conditions. We find that average neural selectivity follows largely the same trend as in the case of normal aging (Fig. 9). This suggests that networks with time-varying learning rates can be mapped to networks with constant learning rates by adjusting the length of training loops during these periods. The implication for our results is that we can interpret the development phase of our network as functionally equivalent to a critical learning period with higher rates and a shorter learning interval compared to the aging phase.

**Figure 9:**
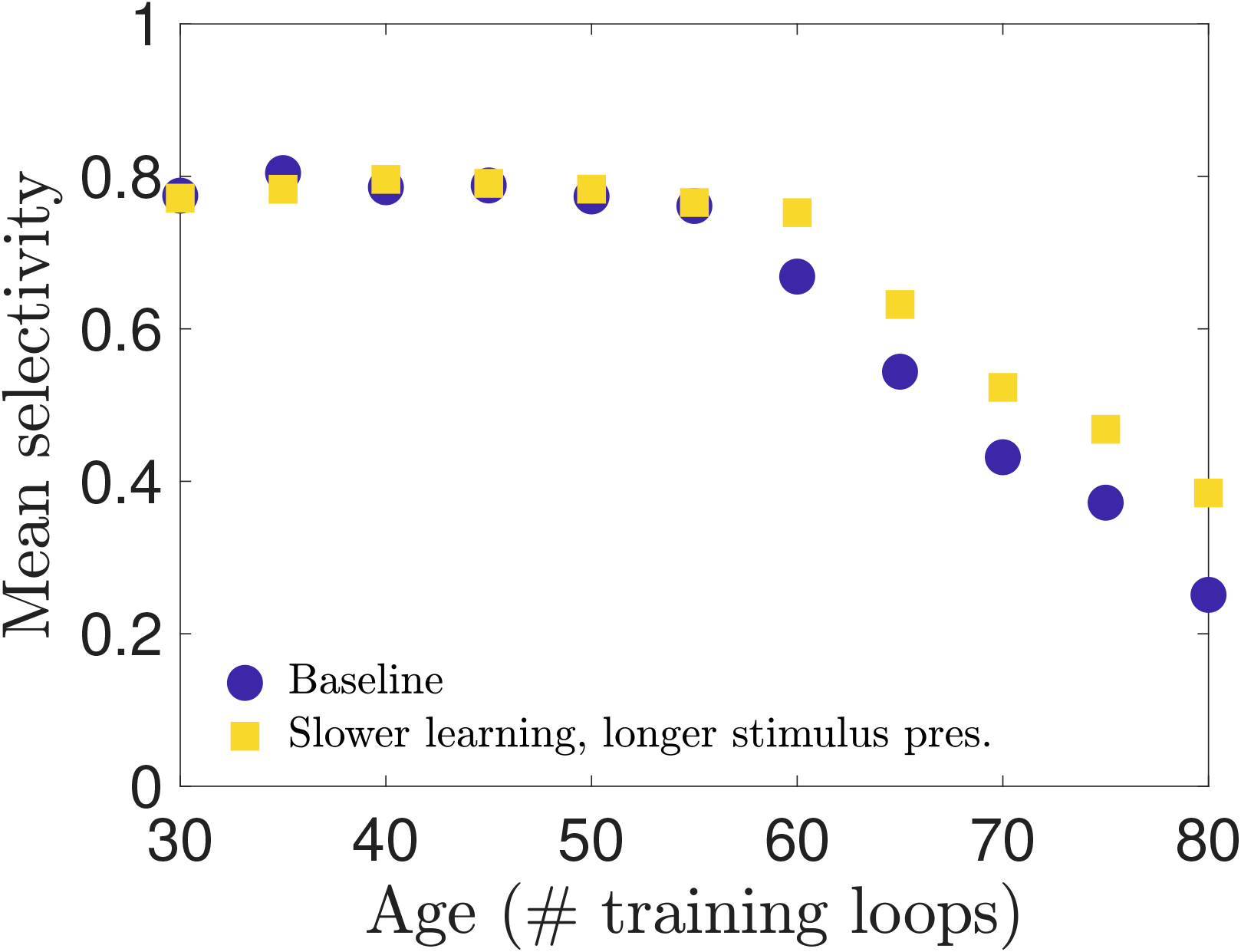
Testing selectivity in a network with a critical learning period: To show that one could in principle implement a critical learning period in the training phase of our network model, we show that if we train a new network in which learning rates are reduced by a factor of 10 and duration of training increased by 10 (representing the end of the critical learning period), the results are qualitatively similar. Thus, the training procedure employed in our model is expected to produce approximately the same results as a model with a critical period during which learning rates vary nonlinearly during training, so long as those changes are compensated by reciprocal changes in the duration of training.

#### Excitatory neurons with similar receptive fields effectively mutually inhibit each other

To gauge the extent to which excitatory neurons with similar receptive fields inhibit one another, we take element (*i,j*) of the product of the lateral connection matrices *W_EI_* and *W_IE_* to approximate the net charge transfer capacity from pre-synaptic excitatory neuron *i* to post-synaptic excitatory neuron *j*, by way of all disynaptic pathways through inhibitory neurons. (Recall that in E-I Net there are no direct E-to-E connections; i.e., *W_EE_* = 0). The greater the magnitude of (*W_EI_W_IE_*)_*ij*_, the greater the inhibition of excitatory neuron *i* by excitatory neuron *j*. As in Fig. 6B of [39], we plot the RF overlap, computed as the cosine similarity of the vectorized input weight matrices, against the elements of |*W_EI_W_IE_*| for all ordered pairs of excitatory neurons. In Fig. 10 we see, similar to [39], that a larger overlap is generally associated with a larger in magnitude *W_EI_W_IE_* value and thus, per our interpretation, stronger inhibition of one excitatory neuron in the pair by the other.

**Figure 10:**
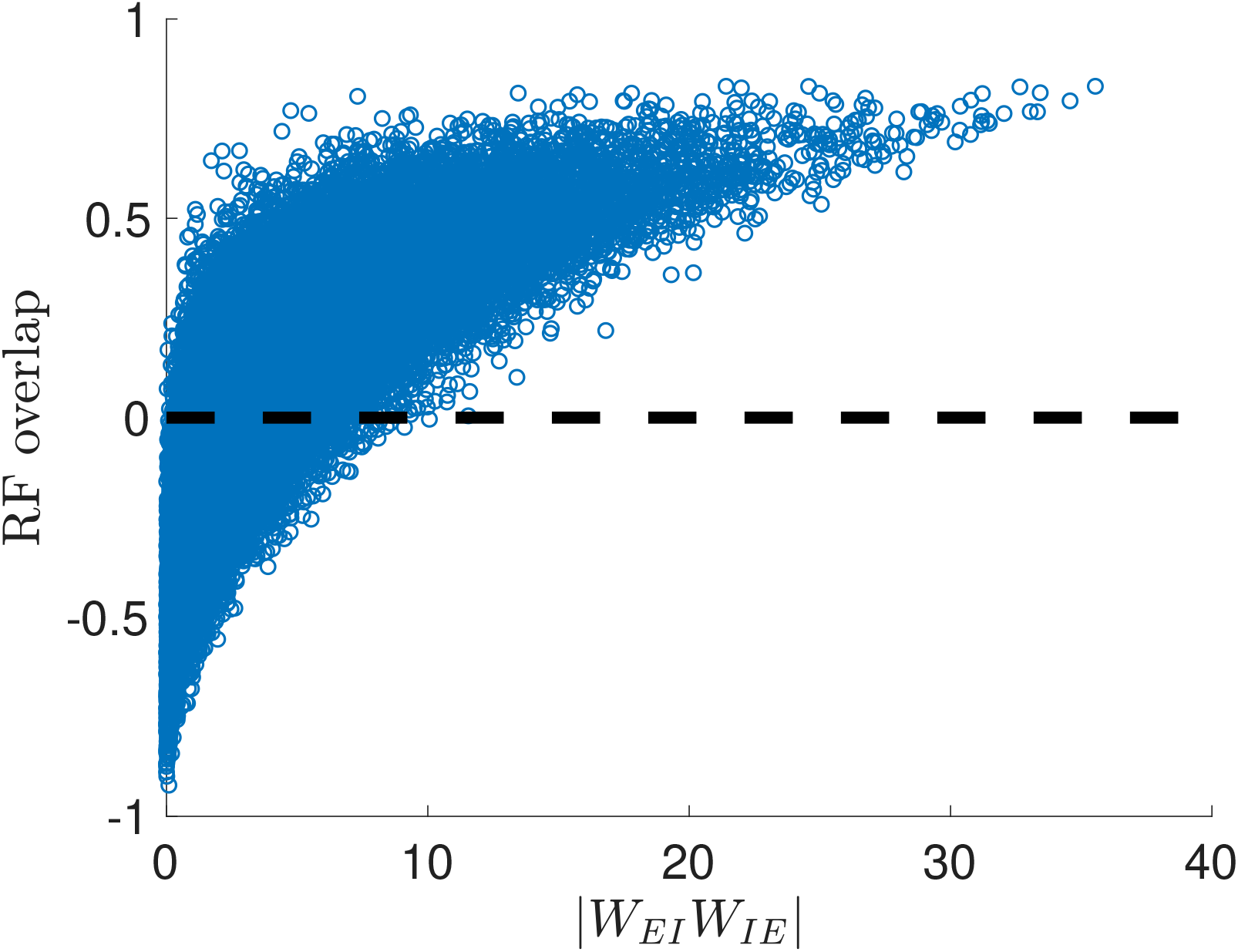
Excitatory neurons with similar receptive fields (RFs) effectively inhibit one another: As a proxy for the total amount of charge excitatory neurons transmit between each other, we use the elements of the matrix *W_EI_W_IE_*, which correspond to the effective weights of the disynaptic connections linking excitatory neurons through single inhibitory interneurons. (There are no direct E-to-E connections in our network). We plot the magnitude of (*W_EI_W_IE_*)_*ij*_ for each ordered pair of neurons *i* ≠ *j* against the overlap of the receptive fields of those same two neurons. As seen in the plot, neurons with the largest magnitude of inhibitory charge transfer tend to have relatively large receptive field overlaps; i.e., excitatory neurons that effectively inhibit each other most strongly tend to have very similar RFs.

#### State dependence of the network selectivity as a function of the target spike rate *p_E_*(*t*_loop_)

We also check the robustness of our results and potential history dependence of the increase in *p_E_*(*t*_loop_) by training the model with different initializations. In Fig. 11A we plot the mean network selectivity obtained by initializing the target excitatory spike rate at *p_E_*(0) = 0.01 (the baseline network used throughout the main text), 0.05, 0.07, and 0.09. We find that although the trajectories appear different, the mean selectivities are comparable at ages for which the networks have similar values of *p_E_*(*t*_loop_). For example, the selectivity when *p_E_*(*t*_loop_) = 0.07 at age 60 in the baseline network is similar to the young selectivity in the network for which *p_E_*(0) = 0. 07. We demonstrate this directly by plotting the mean selectivity versus pE for each network, shown in Fig. 11B. This suggests the network is approximately a state function of the excitatory target firing rate, with a weak dependence on the training history. Moreover, the results in Fig. 11 suggest a lower limit of the mean selectivity of these networks, saturating at a lower bound of approximately 0.2.

**Figure 11:**
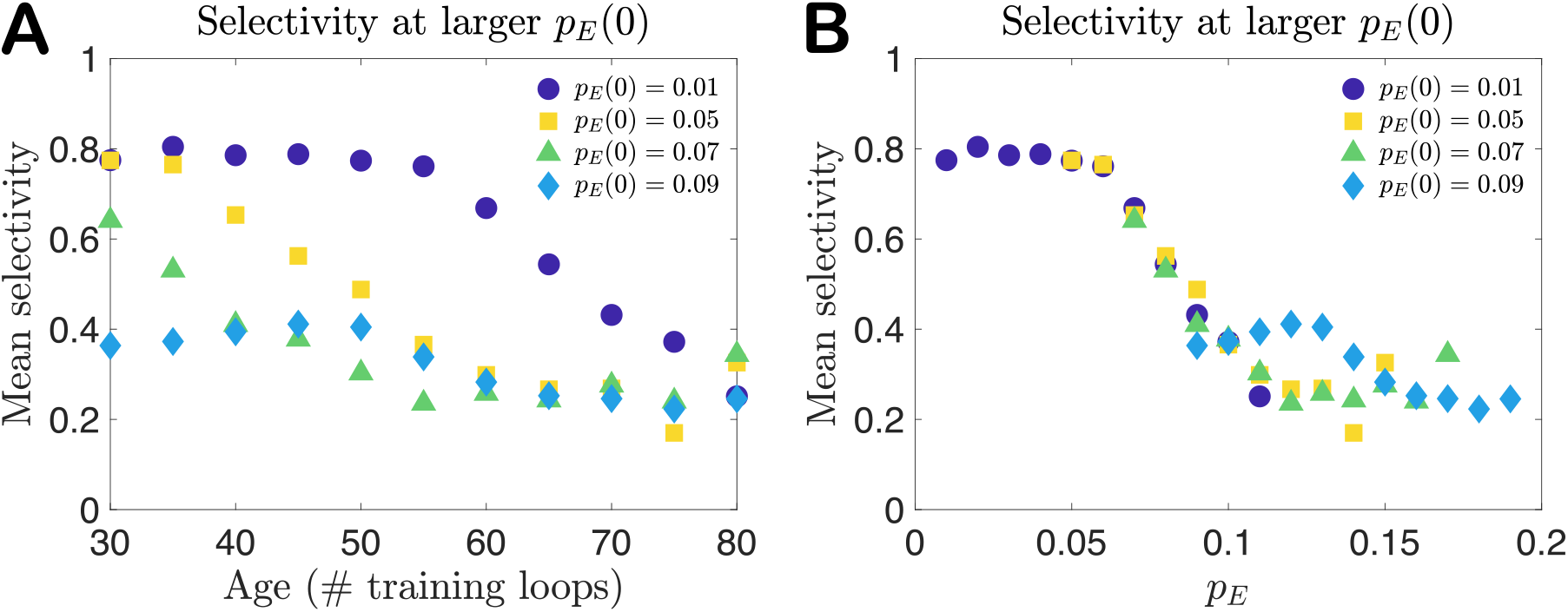
Mean selectivity for larger initial target firing rate: **A**. If the initial excitatory target spike rate is set to a larger value than the baseline value of *p_E_*(0) = 0.01 used in the main text, the mean selectivity that develops in maturity (30 loops) is comparable to that of the baseline network when *p_E_*(*t*_loop_) reaches that same value. For example, the selectivity when *p_E_*(*t*_loop_) = 0.07 (at age 60) in the baseline network is similar to the mature selectivity in the network for which *p_E_*(0) = 0.07. This appears to hold up to values around 0.09, at which the mean selectivity seems to have reached a lower limit of around 0.2. **B**. To demonstrate this more clearly, we plot the mean selectivity versus *p_E_* for each network (using the same data shown in **A**), showing that when the different networks have the same value of pE they have similar mean selectivities. These results suggest the selectivity of a network is approximately a state function of *p_E_*(*t*_loop_).

## Figures

Figures created in MATLAB were saved using the `export_fig` script, available on GitHub [78].

## Supplementary Information

We include a series of supplementary figures to support the findings presented in the main work. Fig. 12 displays histograms of the trained network parameters *Q* (input weights), *W* (lateral weights), and *θ* (firing thresholds) at several intermediate ages following network maturity (30 training loops). This expands on Fig. 2 in the main text.

**Figure 12:**
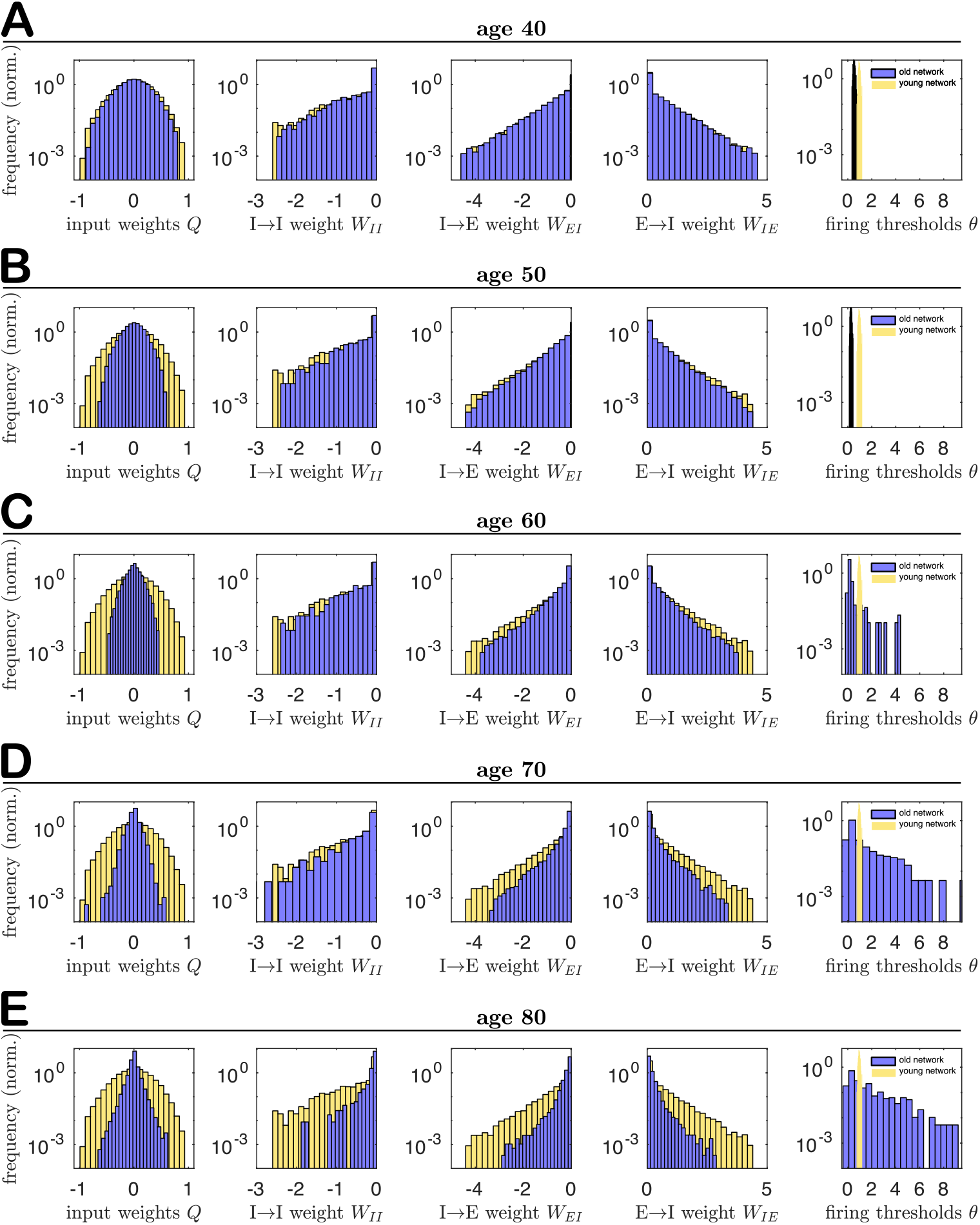
Supplementary Figure: Histograms of physiological network parameters during the aging process: The empirical distributions of the input weights *Q*, lateral weights *W*, and firing thresholds *θ* at different ages during the aging process. **A**. 40 loops, **B**. 50 loops, **C**. 60 loops, **D**. 70 loops, and **E**. 80 loops (same data shown in Fig. 2). This network was trained on frames from movie¤l in the CatCam database [66].

We test the robustness of our results by training the network on three different movies from the CatCam repository (`movie01.tar, movie07.tar`, and `movie16.tar`) and resetting all random number generation before each run. We find no appreciable differences across the results on the three image sets. See Supplementary Figures 13-19.

### Physiological parameter distributions during the aging process

### Physiological parameter distributions

**Figure 13:**
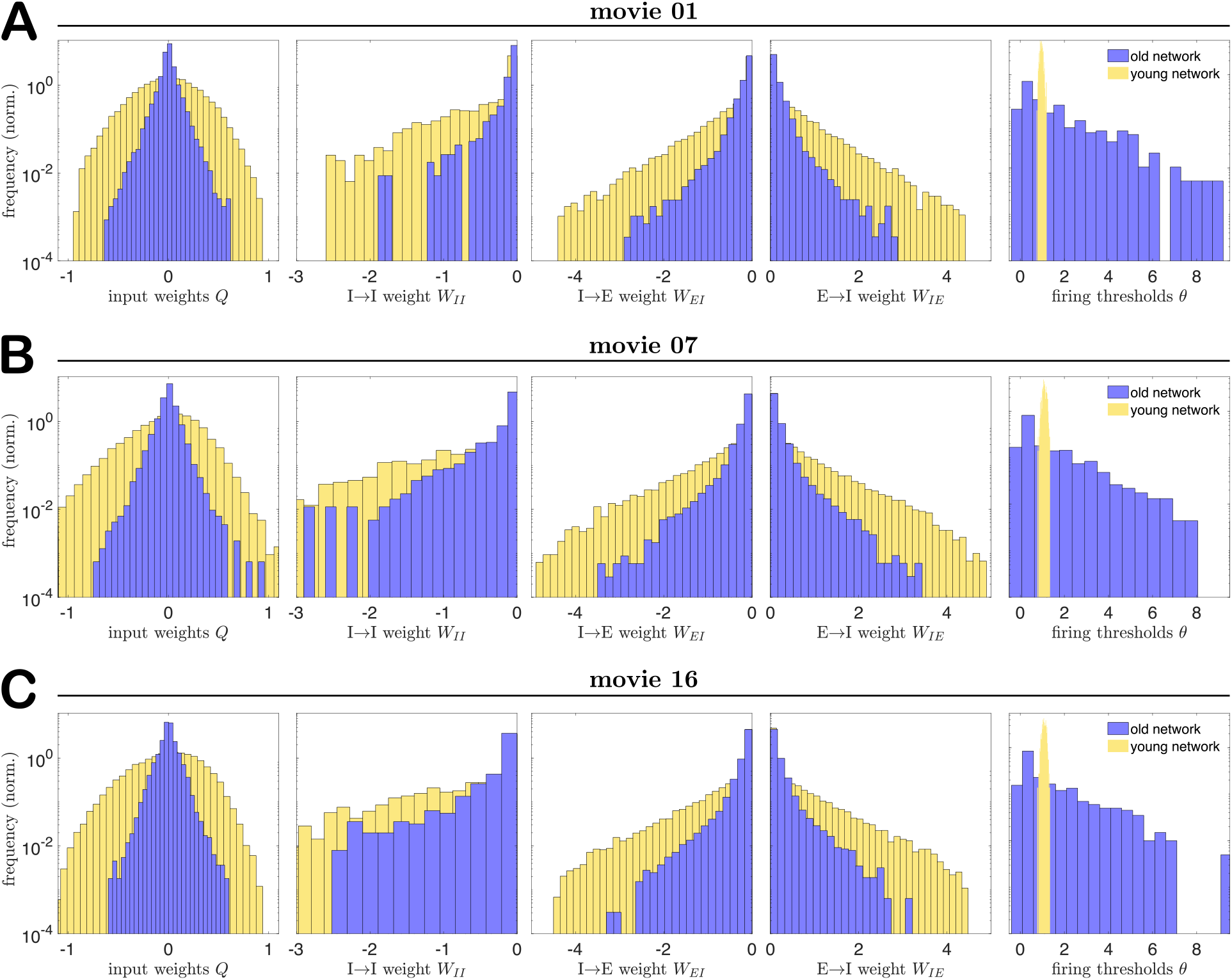
Supplementary Figure: Histograms of physiological network parameters in youth vs. old age for different training movies: The empirical distributions of the input weights *Q*, lateral weights *W*, and firing thresholds *θ* obtained in networks trained on **A**. movie01 (same as Fig. 2), **B**. movie07, and **C**. movie16 from the CatCam database [66].

### Distributions of angles between receptive fields with age

**Figure 14:**
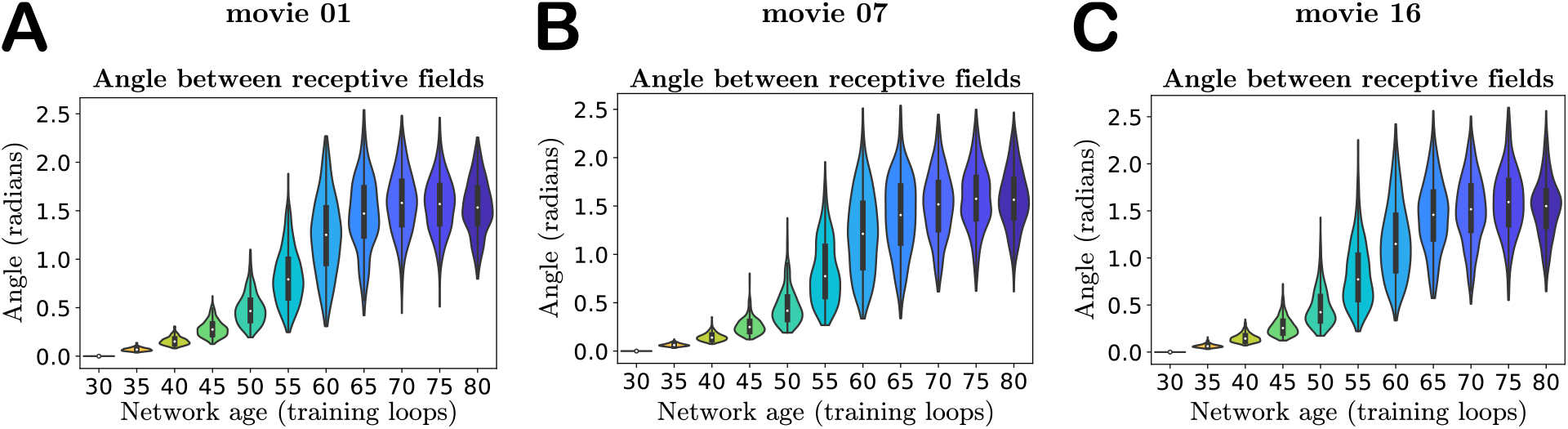
Supplementary Figure: Angles between young and old receptive fields in the input weight parameter space: The distribution of angles between young and old receptive fields in networks trained on **A**. `movie01` (same as Fig. 3B), `movie07` (panel **B**), and `moviel6` (panel **C**) from the CatCam database [66].

### Spike count distributions

**Figure 15:**
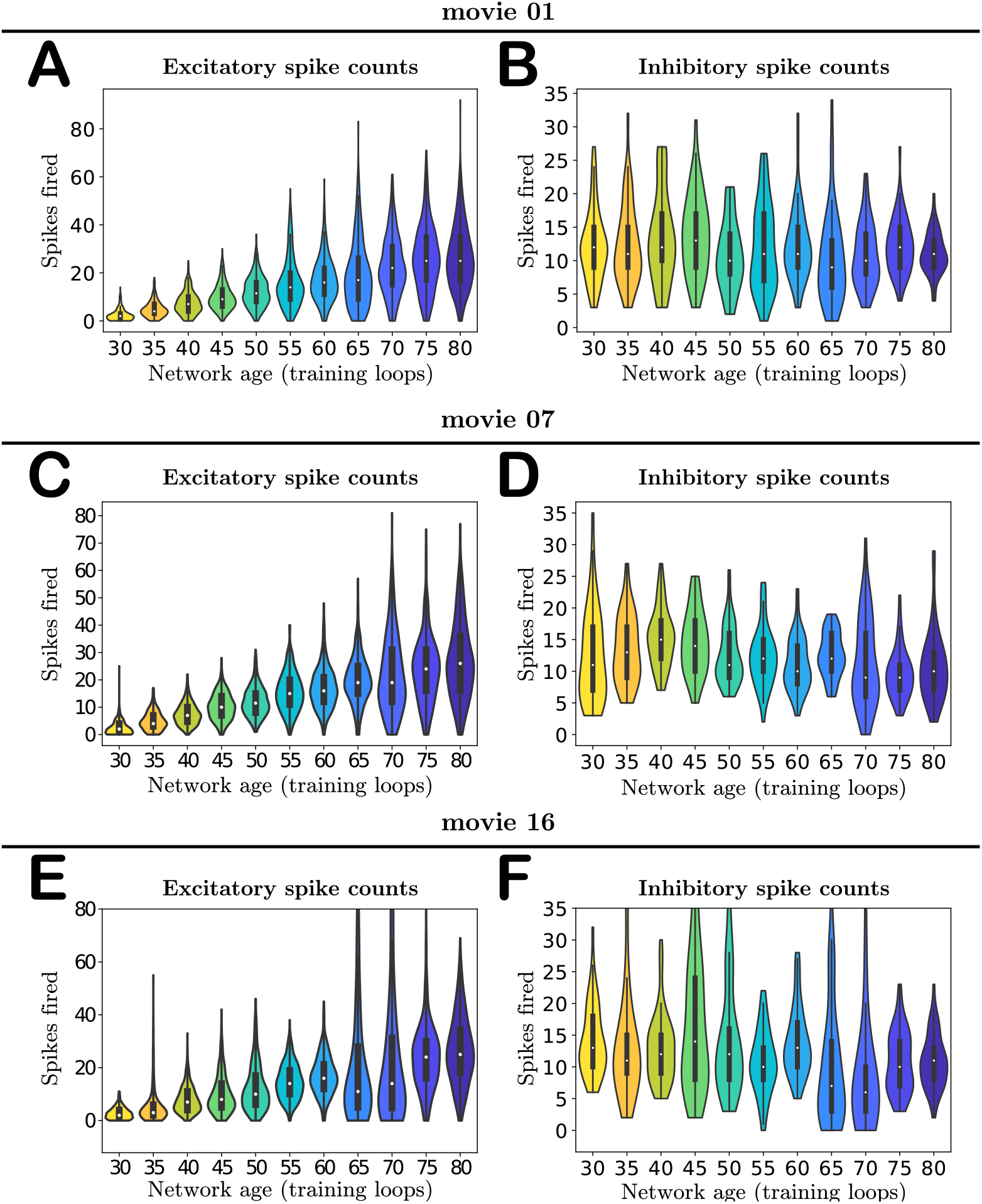
Supplementary Figure: Activity of the network with age: The spike counts of neurons for networks trained on `movie01` (panels **A** and **B**, same as Fig. 4), `movie07` (panels **C** and **D**), and `movie16` (panels **E** and **F**) from the CatCam database [66]. In `movie16` there were some outlier spike counts (e.g., the largest excitatory spike count was 243 at loop 65), so we have restricted the vertical axis to match the range observed for the other movies.

### Orientation selectivity cumulative distribution functions

**Figure 16:**
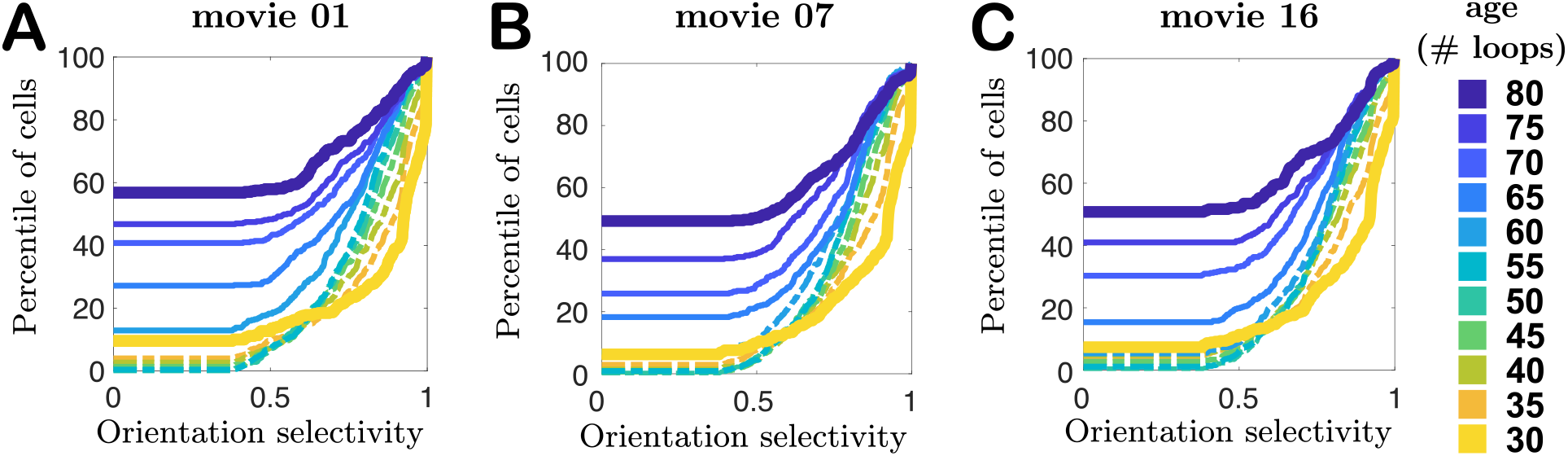
Supplementary Figure: Orientation selectivity changes: The empirical cumulative distribution functions (CDFs) for the orientation selectivity measured in networks trained on images from **A**. `movie01` (same as panel **B** in Fig. 5), **B**. `movie07`, and **C**. `movie16` from the CatCam database [66].

### Orientation selectivity in networks with frozen young parameters

**Figure 17:**
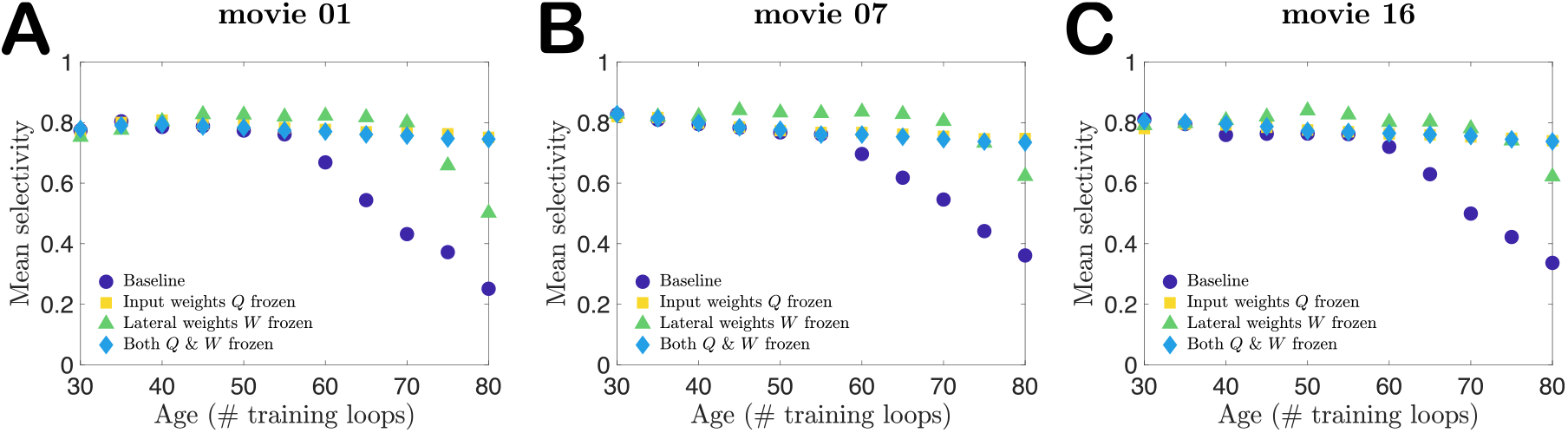
Supplementary Figure: Selectivity of cells when connection weights are frozen during training: The mean orientation selectivity across neurons for networks trained on images from **A**. `movie01` (same as Fig. 6), **B**. `movie07`, and **C**. `movie16` from the CatCam database [66].

### Orientation selectivity in networks with remapped input weights

**Figure 18:**
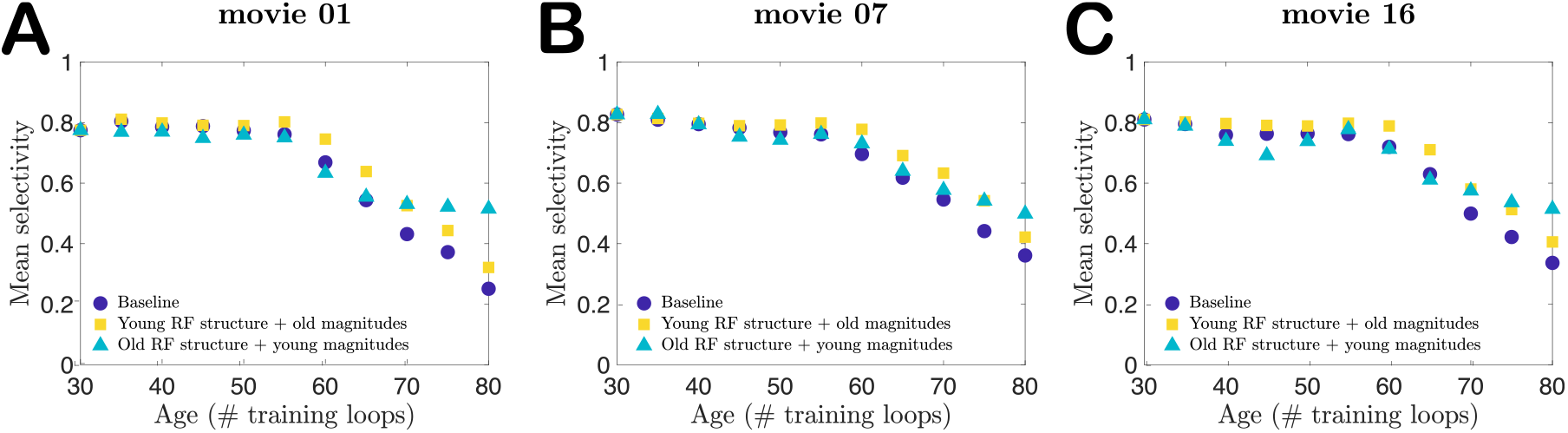
Supplementary Figure: Selectivity of cells with remapped receptive fields: The mean orientation selectivity across neurons for networks trained on images from **A**. `movie01` (same as Fig. 7), **B**. `movie07`, and **C**. `movie16` from the CatCam database [66].

### Parameter swapping numerical experiments

**Figure 19:**
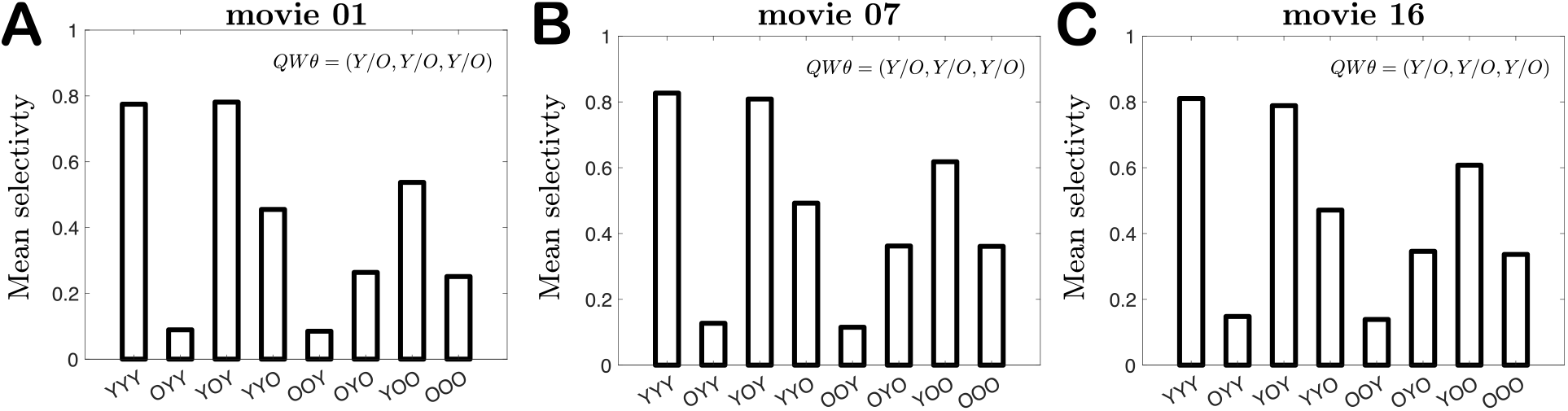
Supplementary Figure: Parameter swap tests: Bar charts of the mean selectivity of several hybrid networks created by swapping the young (Y; age 30) input weights *Q*, lateral weights *W*, or thresholds *θ* with their older (O; age 80) counterparts. The choice of parameters is indicated by the horizontal axis labels. Each bar chart corresponds to a network trained on a different movie from the CatCam database [66]: **A**. `movie01` (same data as Fig. 8), **B**. `movie07`, and **C**. `movie16`.

